# Environmental sensing in dynamic quorum responses

**DOI:** 10.1101/745091

**Authors:** Eric K. Chu, Alex Groisman, Andre Levchenko

## Abstract

Cell communication and coordinated cell behavior are hallmarks of multicellular behavior of living systems. However, in many cases, including the ancient and archetypal example of bacterial quorum sensing, the meaning of the communicated information remains a subject of debate. It is commonly assumed that quorum sensing encodes the information on the current state of the colony, including cell density and physical colony confinement. Here, we show that quorum sensing can also be exquisitely sensitive to dynamic changes in the environment, including fluctuations of the prevailing nutrient source. We propose a new signaling mechanism accounting for this sensory capability. This mechanism combines regulation by the commonly studied *lux* operon-encoded network with the environmentally determined balance of protein synthesis and dilution rates, dependent on the rate of cell proliferation. This regulatory mechanism accounts for observed complex spatial distribution of quorum responses, and emergence of sophisticated processing of dynamic inputs, including temporal thresholds and persistent partial induction following a transient change in the environmental conditions. We further show that, in this context, cell communication in quorum sensing acquires a new meaning: education of cells within a population about the past history of transient exposure to adverse conditions by a subset of induced cells. In combination, these signaling and communication features may endow a cell population with increased fitness in diverse fluctuating environments.

## Introduction

Collective cell behaviors coordinated by inter-cellular communication are ubiquitous across all domains of life^1-5^, supporting evolutionary advantages of multicellular vs. cell-autonomous signal detection and responses. However, the information conveyed in cell-cell communication and its consequences for the population fitness in dynamically changing environments are frequently not known. Quorum sensing (QS) is an ancient archetypal cell-cell communication mechanism found in many prokaryotes^6^. It is used to coordinate collective cell responses, through endogenously-produced diffusible molecules called autoinducers (AI)^7,8^. What exactly is sensed in the process of QS, however, is a matter of debate. Classically, QS has been defined as a means to condition adaptive responses on achieving sufficient cell density (CD), which is gauged by the accumulation of AI with increasing cell number^9,10^. However, AI can also accumulate to high levels due to poor diffusive characteristics of the local environment, even if the cell density is relatively low^11,12^. This observation led to an alternative interpretation of AI-mediated QS signaling as an active method of surveying the diffusive (DF) transport characteristics of the extracellular milieu^13^. Importantly, QS is frequently coupled to stress-response phenotypes such as sporulation^14,15^, virulence^16-18^, and biofilm formation^19-21^, indicating that at least some aspects of QS can report on adverse environmental conditions. Indeed, poor local diffusive properties or high cell density might also hamper nutrient availability^22^; however, neither CD nor DF sensing interpretations directly relate to the detection of changes in the environmental conditions, particularly of subtle fluctuations in nutrient content. It is not clear, therefore, whether QS might in fact be a mechanism to sense dynamic changes in the potentially stressful, dynamic cell microenvironment^23^, or it just enables this sensing (e.g., in a cell density-dependent fashion) through alternative sensory mechanisms^24^.

To explore the function of QS, it is important to investigate the dynamic features of QS responses. However, the emphasis has traditionally been placed on the categorical on or off description, stemming from the common feature of the molecular circuits underlying QS – the positive feedback. Positive autoregulation is observed, for example, in a frequently studied QS circuit of the marine bacterium *V. fischeri*, whose response is controlled by the *lux* operon^25^. It involves AI, produced by the AI synthase LuxI, binding to its cognate cytoplasmic AI receptor LuxR, and positively auto-regulating the *LuxI* and *LuxR* in addition to driving bioluminescence gene transcription^26-28^. This simple feedback regulatory circuit enables a switch-like increase in the QS response after the threshold concentration of AI is exceeded^29^. However, the time scale of this response and its single-cell properties remain poorly understood. It is therefore not clear how the dynamics of QS may compare to other key time scales, including the duration of cell cycle, environmental fluctuations or, for example, in the case of *V. fischeri*, the day-night life cycle of the host (i.e., the Hawaiian Bobtail Squid, *Euprymna scolopes*).

Here, using a highly integrated combination of experimental and computational approaches, relying in particular on a new method to tightly control the coupling between bacterial populations and variable environmental conditions, we found several unexpected properties of QS. In particular, we found that QS cannot be fully accounted for by the current CD and DF sensing theories, thus requiring a new framework for its interpretation. We propose such a framework and provide extensive quantitative evidence for its validity. Furthermore, we find that dynamic, rather than static, QS response can be a critical determinant of the corresponding phenotypic outcomes. In addition, our results suggest that QS can display complex patterns of spatial distribution and variability, allowing for an increased fitness in the uncertain environments, through a communication-driven cell ‘education’ mechanism. These findings can have important consequences for our understanding of QS and the related phenotypic outcomes, and can have implications for other QS-like phenomena, including eukaryotic community effects and other cell communication-related behaviors.

## Results

### QS is defined by a balance between nutrient-defined protein synthesis and dilution due to cell proliferation

To explore whether QS is consistent with either the CD or DF interpretation **(Fig. 1a)**, we studied the QS responses of densely packed colonies of *E. coli* lab strain MG02S ^30^ exposed to various tightly-controlled environments within a microfluidic device. MG02S carries a single, genomically integrated copy of *V. fisheri* LuxIR QS circuit regulating GFP expression in lieu of bioluminescence genes **(Fig. 1b)**. The cells were grown to a completely dense monolayer state within 720 microchambers of identical dimensions, but with variable numbers and sizes of connective channels which modulate the diffusive coupling between the microchambers and the medium-supplying flow-through channels **(Fig. 1c, e, Fig. S1a, b)**. We found that, despite achieving dense colonies of ∼4000 cells each that were grown chemostatically for 24 hours, cells exhibited no detectable QS in the presence of glucose-rich (20 mM) tryptone-complemented medium (TCM) at 25°C **(Fig. S2a, Movie 1)**. However, switching to glucose-deficient TCM elicited a strong auto-induced QS response, consistent with previous reports of glucose-mediated repression of the *lux* operon^31,32^. Furthermore, despite the same cell density and number within each chamber, the magnitude of QS responses across chambers varied for different coupling configurations **(Fig. 1d, S3)**. Thus, the QS responses was not fully consistent with the CD interpretation, but might instead have reflected DF sensing, in which QS response is defined by the diffusion-limited local accumulation of AI. We indeed confirmed that different environmental coupling of individual chambers resulted in variable transport **(Fig. S1c, d, Movie 2, 3)** and cell growth rates **(Fig. 1e, Fig. S1e-g, Movie 4)**. The transport and cell growth were closely related to each other **(Fig. S1h)**, suggesting that diffusive coupling controlled both AI accumulation and cell proliferation via nutrient access. However, contrary to what was expected from the DF sensing interpretation, the magnitude of QS induction (at 24 hours following the medium switch) was not a monotonic decreasing function, but rather exhibited a biphasic dependence on cell growth rate (and diffusive coupling), **(Fig. 1f, i, Movie 5)**, with the response maximized at intermediate growth rates.

**Figure 1.**
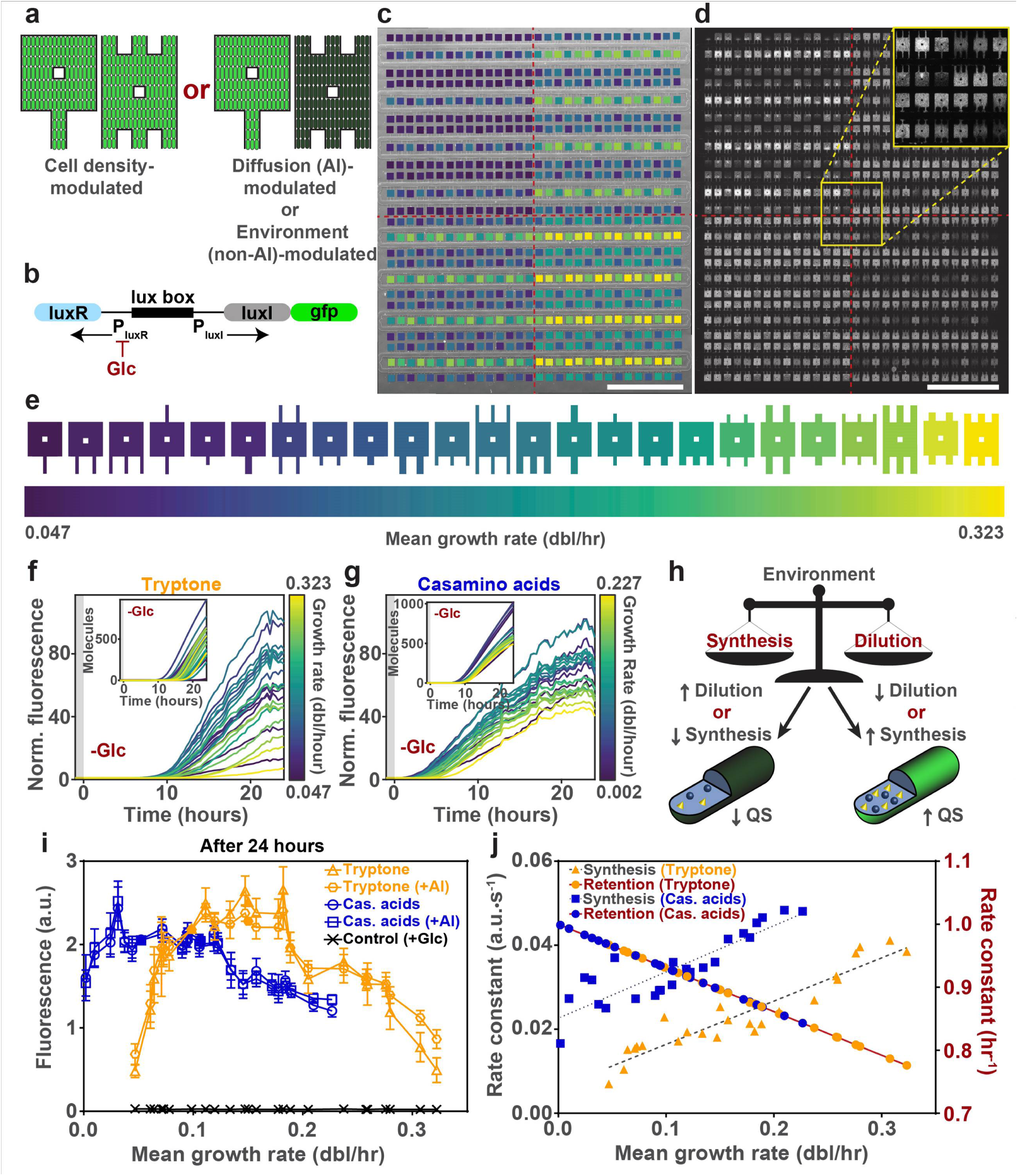
QS is defined by a balance between nutrient-defined protein synthesis and dilution due to cell proliferation. **a**, QS is traditionally interpreted as either a mechanism for cell number/density sensing or diffusive environment sensing, which would lead to different response outcomes in certain environments. **b**, Diagram of QS circuit used in the study. **c**, Phase contrast micrograph of microfluidic device chamber array overlaid with color-coding corresponding to the 24 unique chamber configurations. Scale bar, 1 mm. **d**, Fluorescence micrograph of microfluidic device chamber array taken 24 hours after auto-induced QS response in tryptone medium at 25°C. Inset contain magnified view of the region indicated, which contains all 24 chamber configurations. Scale bar, 1 mm. **e**, Color-coding of chamber configurations based on mean growth rate in tryptone medium. **f, g**, Auto-induction response dynamics over 24 hours in tryptone **(f)** and casamino acids **(g)** media. (mean, n = 6, from 3 independent experiments). Insets denote simulated dynamics obtained by varying only synthesis and dilution with parameters from **(j). h**, Proposed regulatory mechanism of QS response based on the balance of synthesis and dilution of QS machinery proteins. **i**, Distributions of QS response magnitudes after 24 hours in various nutrient and induction conditions. Filled shapes indicate the 2 chamber types excluded from this analysis due to partial formation of cell bilayer (Supp. Info.) (n=105 or n=60, for single- or double-sided chambers, respectively, from 3 independent experiments, mean ± SD). **j**, Measured synthesis and dilution (growth) rate constants from tryptone and casamino acids media conditions.

Further, to directly test whether differential AI accumulation between chambers was responsible for the different QS responses, as would be predicted by the DF sensing hypothesis, we repeated the experiment in the presence of 1 µM of exogenous AI, which is expected to saturate the circuit and thus result in uniform QS induction across different chambers. Unexpectedly, induction with exogenous AI again resulted in a biphasic QS induction pattern, essentially identical to that resulting from auto-induction, albeit with different kinetics **(Fig. 1i, Fig. S2b, Movie 6)**, suggesting that neither CD nor DF sensing hypothesis was sufficient to explain the observed responses.

Since QS response varied with cell growth rate, we tested whether it would change with another source of nutrients. Strikingly, replacing tryptone with a very similar nutrient source, casamino acids (tryptone results from partial and casamino acids from complete digestions of casein, respectively), resulted in a very distinct, more homogeneous response distribution, albeit still biphasic in shape, for both auto- and exogenous AI induction conditions **(Fig. 1i)**. Even more surprising, given the similarity of the nutrients, was the dramatically different dynamics of the QS response in the presence of TCM vs. casamino acid complemented medium (CACM) (**Fig. 1g, Fig. S2c, S4a, b, Movie 7-9)**.

Our results suggest that neither CD nor DF interpretation is completely adequate to account for the observed QS responses, with the growth rate being a more predictive determinant of the response than either cell density or local accumulation of AI. An explanation of this growth dependency might be provided by the dynamics of QS response, and in particular, the observation that the time scales of QS are comparable to and can exceed the duration of the cell cycle. Therefore, the dynamics and amplitude of the response can be strongly influenced not only by the biochemical interactions within the molecular QS circuit, as is commonly assumed, but also by the dynamic balance between protein synthesis and growth-mediated protein dilution **(Fig. 1h)**. We indeed found that the synthesis and dilution (which can also be converted to protein retention, as shown here) of QS-regulated protein (GFP) were monotonically increasing functions of the cell growth rate **(Fig. 1j)** for both tryptone and casamino acids as nutrient sources. However, the average growth rate was higher and synthesis rate was lower for the tryptone vs. casamino acids complemented media. To test whether the balance of QS protein synthesis and dilution would be able to account for various experimentally observed response dynamics, we modified previously published mathematical models of QS response^33,34^ to incorporate the measured synthesis and dilution parameters from the different conditions **(Fig. S5)**. By matching the synthesis and dilution rate constants to the experimentally estimated values, we obtained the predicted QS dynamics that was consistent with the experimental observations for both nutrient sources **(Fig. 1f, g, Fig. S6)**. This result provided first evidence for the key influence of the balance between protein synthesis and dilution as a regulatory mechanism of the environmental sensitivity of the QS response.

### Information about distinct environmental conditions is reflected in QS response dynamics

To further support the protein synthesis/dilution balance (PSD) mechanism, we explored QS under additional environmental perturbations. In particular, we reasoned that increasing the temperature would accelerate cell growth and thus increase protein dilution, leading to an attenuated QS response. Indeed, increasing the temperature from 25°C to 30°C elevated cellular growth rates resulting in a complete abrogation of response in TCM even after 48 hours of observation **(Fig. 2a,b, Fig. S2d, Movie 10)**. The same temperature increase in CACM, which has reduced growth rates, resulted in a partial QS onset (88 out of 144 chambers assayed) that was delayed by approximately 15-20 hours, matching the mathematical model predictions **(Fig. S2e, Movie 11)**. To directly test whether protein abundance could modulate the onset of the QS response, we transformed the cells with a construct coding for IPTG-inducible *LuxR* expression **(Fig. S4c)**, and used it to examine the effect of LuxR overexpression on QS induced in TCM at 30°C. Overexpression of LuxR resulted in the rescue and immediate onset of QS response in all chambers even at the elevated temperature **(Fig. 2c, Fig. S2f, Movie 12)**. As expected, the timing of the onset of signaling was also dramatically shortened with overexpression of LuxR in TCM at 25°C, with responses in all chambers triggered virtually immediately **(Fig. S2g, Movie 13)**. Alternatively, if the level of glucose used for cell culture prior to the switch to TCM was lowered (20 to 10 mM), which was expected to increase the basal level of LuxR expression due to both partial catabolic de-repression and reduced cell growth, a decrease in the QS onset timing and increase in the amplitude was observed **(Fig. S2h, Movie 14)**. Although these experiments suggested the importance of achieving a high level of LuxR expression for QS onset, the onset time was also dramatically decreased if exogenous AI was added at the time of the QS induction in TCM and CACM at 25°C **(Fig. S2b, c)**. This result implied that QS onset can be accelerated if the abundance of either LuxR or LuxI (and thus AI) is increased in the cells, which can be controlled by the balance of synthesis and dilution of these proteins, as dictated by growth conditions. On the other hand, consistent with the results above, the response magnitude, unlike the response dynamics, was insensitive to exogenous AI (**Fig. 1i, Fig. S4d)**, implying that AI, under these conditions primarily controls the dynamics of the QS responses (Fig 2d). All of these results were also well approximated by the corresponding versions of the mathematical model, further supporting the PSD mechanism **(Fig. S2, 6)**.

**Figure 2.**
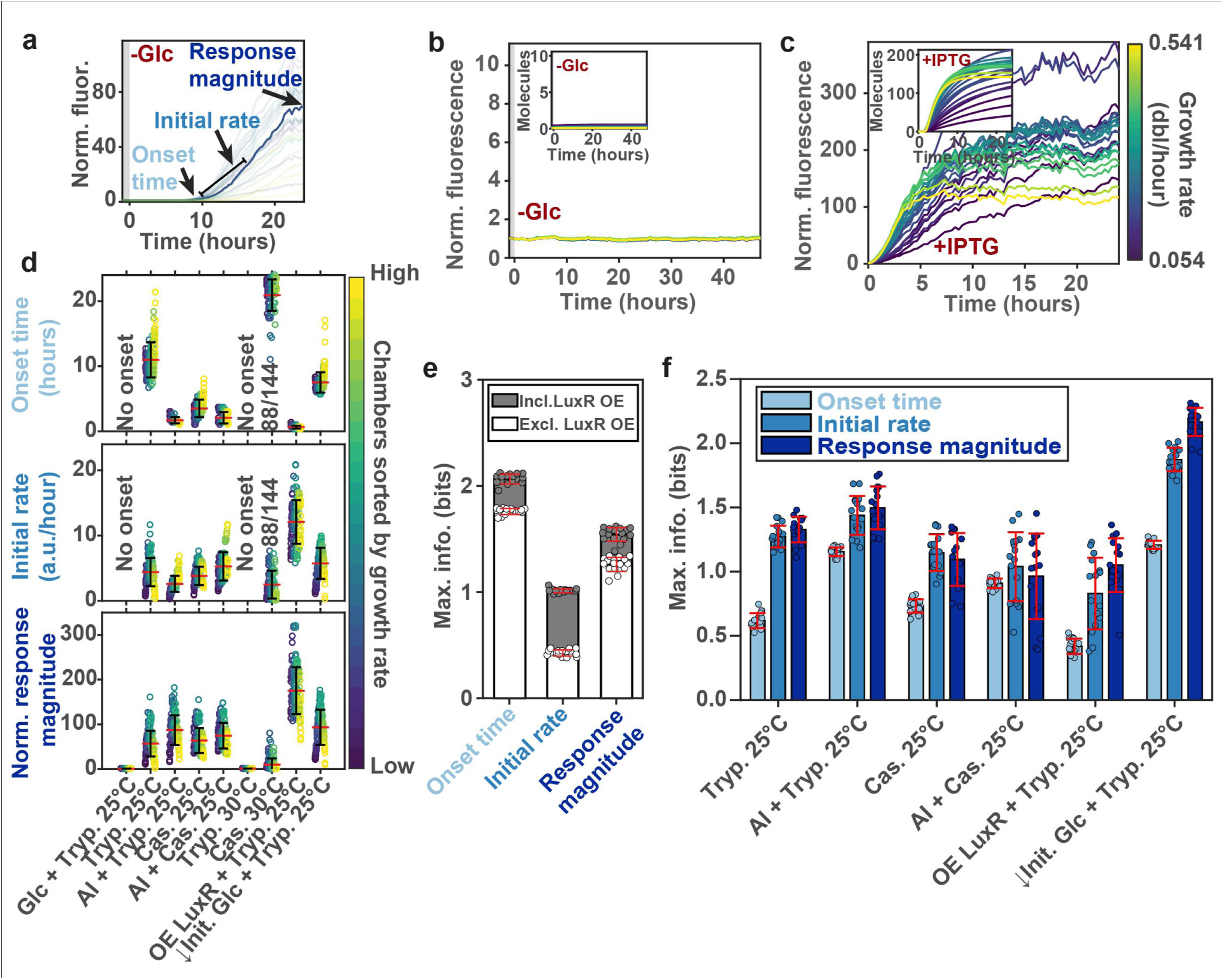
Information about distinct environmental conditions is reflected in QS response dynamics. **a**, Separation of QS dynamics into three components: onset time, initial response rate and 24^th^ hour response magnitude. **b,c**, Response dynamics within, **(b)** tryptone medium at 30°C, and **(c)** tryptone medium with 1 mM IPTG (for LuxR overexpression) at 30°C (mean, n = 6, from 3 independent experiments). Insets denote simulated dynamics. **d**, Summary of onset times, initial rates, and response magnitudes from various conditions (n = 144, from 3 independent experiments). **e**, Comparison of information about global environmental conditions discerned by each component of QS dynamics assuming all chamber configurations are equivalent. **f**, Comparison of information about the local differential coupling environment discerned by each component of QS dynamics for various global environmental conditions. (mean ± SD)

Diverse conditions used in the experiments suggested that the QS outcome can be encoded not only in the magnitude but also in the timing of the onset and the initial rate of the QS response **(Fig. 2a, d, Fig. S6)**. This dynamic information can inform downstream responses long before the steady state QS response is achieved. It is thus of interest to explore the sensitivity of the dynamic aspects of QS to the various environmental conditions and to the degree of coupling between the colony and the environment. One way to assay the sensitivity to environmental inputs is an information theoretic approach^35,36^, which, given the response variability, can indicate how many input types or doses can be accurately resolved. Using this approach, we found that the onset time showed the highest sensitivity in discriminating external environmental conditions explored here, yielding 1.76 ± 0.03 bits of information, equivalent to accurately distinguishing 2^1.76^ = 3.4, or approximately 3 different types of conditions **(Fig. 2d, e, Fig. S7)**. This discrimination capacity was in spite of the variability of environmental coupling across the chambers, suggesting that the timing of the QS onset can provide the early information about the changes in the environment regardless of the exact nature of the coupling of environment to the cellular niche. Conversely, the response magnitude was the metric that was most sensitive to the differential diffusive coupling, with at least 1 bit of information representing the ability to accurately distinguish at least 2 coupling levels for each environmental condition **(Fig. 2d, f, Fig. S8)**. The onset timing, on the other hand, was the least informative about this aspect of the cell environment, consistent with the results above. This result suggested that the degree of cellular confinement in a particular niche can indeed be conveyed by QS, but information about this aspect of cell environment does not become available until high levels of response are attained.

We noted that our calculation of the information about the environmental coupling was likely an underestimation, since the biphasic nature of the QS response distributions meant that two distinct mean colony growth rates could correspond to the same mean QS response magnitude, creating an ambiguity that can reduce the information content in the QS response magnitude. Therefore, relatively slow and relatively fast growth can potentially result in the same QS response magnitude and trigger the same levels of downstream adaptive response, unless there is an additional means to resolve this ambiguity, a possibility that we explored next.

### Spatial properties of environmentally regulated QS response

We hypothesized that the ambiguity associated with the biphasic dependence of the mean QS response on the degree of coupling (growth rate) could potentially be resolved, if two different chambers yielding the same average QS response magnitude at different growth rates would have distinct spatial distributions of the QS signal. In particular, consumption of the nutrients diffusing into the chambers by multiple cells within the chambers can lead to gradients of nutrient availability, and thus, corresponding gradients of the cellular growth rates. As a result, each chamber can span a range of growth rates, with growth rates higher in the chamber regions adjacent to the coupling channels due to proximity to the nutrient source. We thus hypothesized that the range of growth rates within a chamber can result in the range of QS responses corresponding to the biphasic distribution curve measured before **(Fig. 3a)**. Consequently, the spatial gradients of QS response within a chamber would correspond to a segment of the biphasic dependence curve, with the local inclination of the segment specifying the direction of the spatial gradient **(Fig. 3b)**. The biphasic nature of the curve would, therefore, specify gradients of opposite signs, according to the rising and falling parts of the biphasic curve. Since the two values of growth rates corresponding to the same QS response can map to two parts of the biphasic curve with the opposite slopes, the resulting gradients of QS responses would be opposite, providing extra information that can resolve the ambiguity discussed above.

**Figure 3.**
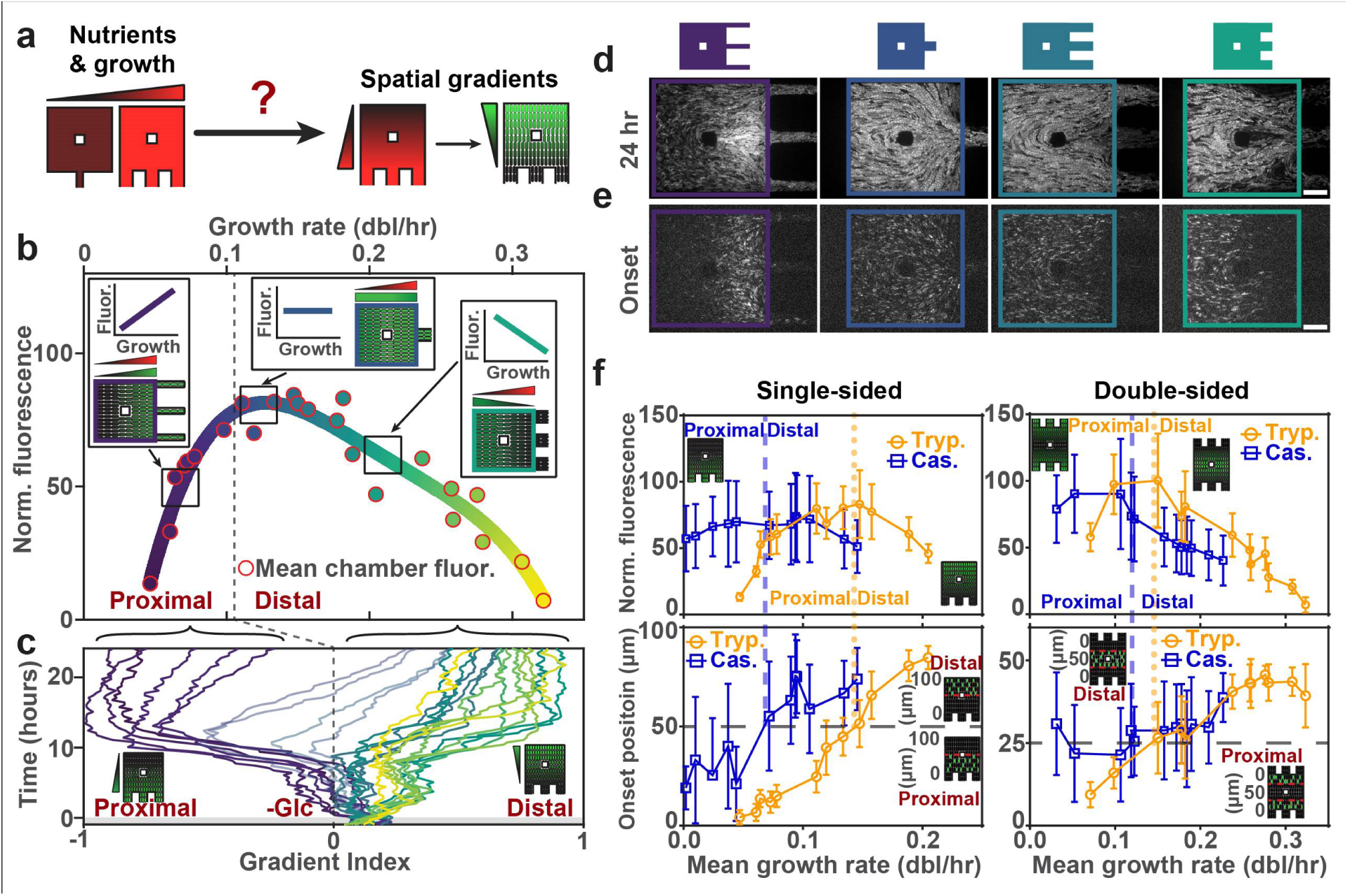
Spatial properties of environmentally regulated QS response dependence. **a**, Spatial dependence could arise from nutrient and growth gradients within the chamber. **b**, The proposed relationship between spatial response gradients and the biphasic response magnitude distribution. **c**, Quantification of auto-induction spatial response gradient dynamics over 24 hours in tryptone medium at 25°C. Positive gradient index indicates higher response in the regions distal from the coupling channels, whereas negative gradient index indicates higher response in the regions proximal from the coupling channels. (mean, n = 6, from 3 independent experiments). **d,e**, Fluorescence micrographs of chamber configurations corresponding to those illustrated in **(b)**, at 24^th^ hour **(d)** and at onset **(e)**. Scale bar, 20 µm. **f**, Spatial localization of auto-induced QS response onset in single- and double-sided chambers in tryptone and casamino acid media at 25°C, in relation to response magnitude distributions after 24 hours in the same conditions (n = 43-64, from 3 independent experiments) (mean ± SD).

We indeed found that different chambers incubated in TCM at 25°C for 24 hours had distinct spatial QS magnitude distributions, and that the spatial gradient directions were fully consistent with the hypothesis above, and thus the PSD hypothesis **(Fig. 3b-d Fig. S9a)**. We further explored if this hypothesis could also account for the dynamics of spatial QS response evolution. In particular, we investigated if this spatial dynamics can be accounted for by the dynamics of the mean QS responses **(Fig. S10, 11)**. For example, the QS response distribution is biphasic throughout the 24 hours of QS induction in TCM at 25°C, implying that the location of maximum QS response in chambers with lower average growth rates would be proximal to the nutrient-supplying channels throughout the response, starting from onset **(Fig. 3c-e, Fig. S10, 11)**. On the other hand, the maximal response would occur in more distal regions for chambers with higher average growth rates. Indeed, we found that the onset position for QS in both TCM and CACM conditions corresponded to the location of the maximal response as well as the local inclination of the biphasic curve, shifting from proximal to distal regions of the chamber when examining chambers with increasing mean growth rates **(Fig. 3f, Fig. S9b, S12)**. Overall, these results suggest that both the average QS response magnitude and the spatial gradients of the QS response in individual chambers can be accounted for by the same mechanism. This observation suggests that the biphasic nature of the response magnitude distribution, explained by the PSD mechanism, can lead to a variety of spatial QS distributions, as a function of coupling between the cellular niches and extracellular milieu.

### Temporal thresholds and bistability in QS response

Our data suggests that the dynamics of QS response can convey the information on a persistent change in the environmental conditions, particularly the nutrient content, with this information further refined as a function of the degree of diffusive coupling, resulting in nonuniform spatial distributions of the signaling magnitude. However, the changes in the environment are frequently transient or display complex dynamics. We thus investigated how QS responds to such dynamic conditions. In particular, we investigated if variable QS onset timing can translate into condition-dependent temporal thresholding, requiring a persistent rather transient change in the environment, longer than a certain threshold, for the response to occur **(Fig. 4a)**. We explored this possibility by transiently switching from the glucose-rich TCM to the QS-inducing TCM at 25°C for 4 hours, which is shorter than the fastest onset timing of ∼7 hours in these conditions **(Fig. 1h)**. This transient change in the environment indeed resulted in no detectable QS response in both simulations and experiments, suggesting a temporal filtering of inputs below a threshold duration **(Fig. 4b, Fig. S13a, Movie 15)**. Furthermore, periodic changes in cell environment between TCM and glucose-rich medium (4 hour-long pulses, 50% duty cycle) again resulted in the absence of QS response, implying no temporal integration of the transient inputs **(Fig. S13b, Movie 16)**. On the other hand, a 16-hour stimulation with the TCM, exceeding the slowest QS onset time among cells, enabled the response in all chambers **(Fig. 4b, Fig. S13c, Movie 17)**. This response recovered to the baseline in all but 3 chambers within 24 hours after restoring glucose-rich conditions. In combination, these results suggested effective temporal filtering of the environmental changes by the QS circuit, ensuring that only sufficiently persistent signals activate the response.

**Figure 4.**
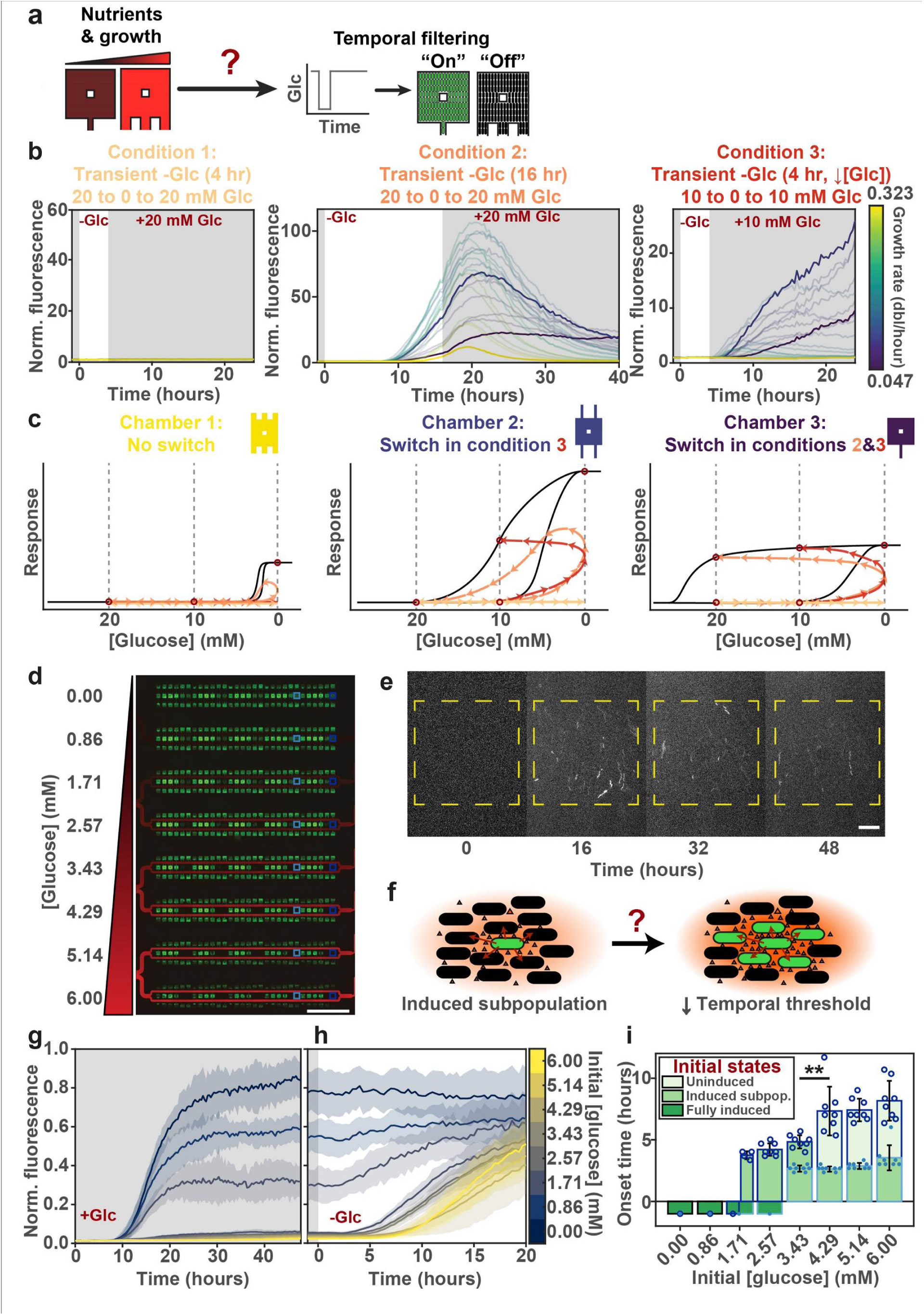
Sensitivity to environmental change as encoded in the form of a temporal threshold can be enhanced with cell signaling. **a**, QS onset can be controlled by a condition-dependent temporal threshold during transient stimulation. **b**, Response dynamics of 4 hour transient glucose (20 mM) removal (condition 1), 16 hour transient glucose (20 mM) removal (condition 2), and 4 hour transient glucose (10 mM) removal (condition 3). All conditions were in tryptone medium at 25°C. (mean, n = 6, from 3 independent experiments). 3 sample chambers in each condition have been highlighted. **c**, Theoretical hysteretic diagrams of three sample chambers within the three conditions in **b. d**, Fluorescence micrograph of gradient-generating device, with chambers of interest indicated in blue boxes and the gradient visualized in red. Scale bar, 1 mm. **e**, Filmstrip of a representative chamber exposed to 2.57 mM glucose demonstrating maintenance of a stably induced subpopulation over ∼40 hours. Scale bar, 20 µm. **f**, Proposed effects of an induced subpopulation. **g**, Response dynamics from chambers exposed to 0 – 6 mM linearly graded glucose concentrations over 48 hours (n = 8). **h**, Response dynamics over 24 hours after glucose removal (n = 8). **i**, Onset time of chambers exposed to different initial concentrations of glucose. (n = 8, \*\**P*<0.01, two tailed Student’s *t*-test). (mean ± SD)

The 3 chambers retaining the QS signaling following transient stimulation were the ones with the slowest growth rates **(Fig. 4b, Fig. S14a)**. Since transient responses can be stabilized by the positive feedback and the associated bistability and hysteresis, we hypothesized that the QS-associated positive feedback might be maintaining the response in a subset of chambers, even in the presence of the glucose-rich environmental conditions. Further analysis and our prior work^30^ suggest that the bistability regime, and thus, the number of chambers retaining the response following transient stimulation, can be expanded by lowering glucose concentrations in the medium prior to and following the transient stimulation **(Fig. 4c)**. Indeed, for cells pre-incubated in 10 mM glucose (mean QS onset time of 7.5 ± 1.5 hours), glucose removal for 4 hours was sufficient to elicit QS in several, but not all, chambers, again supporting the existence of a temporal threshold, albeit lowered for this condition **(Fig. 4b, Fig. S13d, 14b, Movie 18)**. Consistent with the previous results, the chambers showing the QS response in this experiment were also the ones with slower growth rates and thus included those that displayed persistent response after cessation of the stimulus. As hypothesized, the number of chambers where QS was persistently retained after removal of the stimulus increased to 7. Extending the stimulus duration to 9 hours permitted all chambers to become transiently induced and achieve higher amplitudes of induction, but did not affect the number of chambers displaying irreversible QS **(Fig. S13e, Movie 19)**. These results highlighted a strong bi-stable nature of the response consistent with the positive feedback interactions within the *lux* signaling circuit^27,30^ leading to a ‘memory’ of a prior induction even if the environmental conditions no longer favor QS. Overall, we concluded that better environmental coupling and more nutritious environment can impart longer temporal thresholds ensuring a low and transient QS response for a subset of chambers exposed to a transient change in the environment **(Fig. 4c)**. On the other hand, poorer environmental coupling and less nutritious environments can each reduce the temporal threshold and ensure persistent QS responses in an increasing number of micro-chambers **(Fig. 4c)**.

### Cell communication allows a small pre-induced cell sub-population to quicken the onset of response in the rest of the population through cell communication

Visual inspection of chambers with persistent QS response revealed that only small fractions of cells were highly induced, with the rest of the population displaying no detectible QS signaling **(Fig. S14, Movie 17-19)**. The higher-than-basal levels of AI produced by the induced subpopulation could be key to the maintenance of the high stable state of expression of QS genes^37,38^ in the presence of active environmental suppression. Indeed, simultaneous induction and repression with exogenous AI and glucose, respectively, stably maintained an induced subpopulation **(Fig. S15, Movie 20)**. Furthermore, we hypothesized that this subset of cells could carry a memory of a prior stimulation, which can provide a selective advantage if conditions become adverse, and thus, stimulatory again. Although this strategy is superficially similar to the commonly assumed ‘hedging of bets’ scenario^39,40^, which postulates that a diversification of response within a population can confer a selective advantage under uncertain environmental conditions, the cell-cell communication nature of QS imparts additional benefits. In particular, the induced cells can potentially promote induction of neighboring cells after worsening of conditions through secretion of AI, thus quickening the timing of the QS onset in the uninduced cells, permitting a greater fraction of cells to faster assume the required adaptive phenotype. We thus explored if cell communication inherent to QS can indeed reduce the temporal threshold and accelerate the onset of response.

To compare the temporal threshold between fully uninduced and partially induced cell populations, we modified the microfluidic platform to screen distinct environmental conditions that can yield such populations simultaneously. More specifically, we explored 12 chamber configurations in the presence of several glucose concentrations in the 0-6 mM range **(Fig. 4d, Fig. S16)**, which enabled us to elicit a wide spectrum of responses ranging from complete presence to complete absence of induction within the chamber configuration with the highest diffusive coupling **(Fig. S17a, Movie 21)**. We found that in the 0-0.86 mM range of glucose concentrations, the cells in the colonies were fully induced, whereas in the 1.71-6 mM range, we observed bi-stable responses, with chambers showing different fractions of induced cells that were stably maintained for at least 40 hours after initial onset **(Fig. 4e, g, Movie 21)**. In the cases of 1.71 and 2.57 mM, the bi-stability resulted in either complete (ranging from 18.77±8.26% to 97.26±0.72%) or partial (ranging from 0.037±0.066% to 12.19±6.82%) induction in different chambers, while chambers in the 4.29-6 mM range contained either partially induced or completely un-induced (ranging from 0.0015±0.0009% to 0.0079±0.0047%) cell populations **(Fig. S18)**. This range of distinct bi-stable responses allowed us to examine whether and how the presence of different fractions of induced cells would affect the QS response **(Fig. 4f)**.

To accomplish this, we studied the effect of switching to TCM from all glucose supplemented conditions, which resulted in a complete QS induction for all chambers, but with distinct onset times **(Fig. 4h, Fig. S17b, Movie 22, 23)**. In all cases, we compared, for the same glucose level, the onset timing of uninduced cells either in the fully uninduced populations or populations containing a fraction of induced cells, due to the underlying bi-stability. For all glucose levels, we found that the presence of a fraction of induced cells dramatically shortened the QS onset time in uninduced cells **(Fig. 4i, Fig. S19)**. Notably, the most dramatic relative change in the onset timing was observed between 3.43 and 4.29 mM of glucose in the pre-incubation medium, also coinciding with the lowest glucose levels allowing bi-stable responses and thus presence of induced sub-populations. This result was consistent with AI-mediated cell-to-cell communication from the induced subset of cells, pre-conditioning the remaining uninduced cells in the population and thus promoting a quicker QS induction following the onset of adverse conditions. This was also consistent with the faster onset of QS in separate experiments whereby cells were exposed to low doses of exogenous AI **(Fig. S20, Movie 24-26)**. These experiments suggested that if the fitness of the population is coupled to the QS onset, the presence of the persistently induced sub-population can provide a selective advantage to the whole population by enabling a faster response to a dynamic change in the colony environment.

## Discussion

QS is one of the most ancient mechanisms of cell communication, and yet, its functional role remains ambiguous. It has been proposed that, in addition to the widely accepted cell density sensing function, QS can also serve to assess the diffusive transport properties of the microenvironment. Our results provide a more complex view of this archetypal signaling mechanism, demonstrating that QS signaling is also sensitive to dynamic changes of environmental conditions such as nutrient composition and access. The observed QS responses are best explained not only by the currently accepted view of QS as driven by positive feedbacks inherent in most QS genetic circuits, including the *lux* operon explored here, but also by the balance of protein synthesis and dilution modulated by the specific microenvironments that the cells are exposed to. In combination, these regulatory mechanisms endow the QS response with several new properties that can strongly affect the fitness of the cell population and the outcome of the adaptive processes frequently coupled to the QS activation.

We found that a subtle change in the nutrient source, such as the degree of digestion of casein, can strongly influence the balance of the rates of cell growth rate and protein synthesis, translating into the dilution and synthesis of the QS-mediating gene products. In particular, a higher degree of digestion translated into a substantial increase in protein synthesis rate accompanied by a decrease in cell proliferation. The reciprocal balance between protein synthesis and cell proliferation may reflect a more general strategy of allocation of limited available resources to different intracellular processes^41,42^. Importantly, this balance can also be influenced by inputs other than nutrients, including, as shown here, gene amplification or altered temperature. The resulting complex regulation translates into non-monotonic responses of the long term QS amplitude as a function of colony growth rate, which in turn translates into the distinct distributions of the spatial QS responses, with the gradients of the response pointing either toward or away from the nutrient source. This inherent non-monotonicity might also explain complex differentiation patterns of *B. subtilis* into a state of competence^43^, suggesting similar regulation in a distinct underlying QS circuit.

Successful adaption may depend not only on mounting the responses that are adequate to the environmental challenges, but also on how quickly these responses can be mounted. The bistability inherent in the positive feedback regulation of most known genetic QS circuits, when modulated by the environmentally defined variable balance between growth rate and protein synthesis, allows for complex decision-making. In particular, this regulatory mechanism can ignore pulsatile environmental inputs, such as changes in nutrient content, that are too transient, while responding to more persistent stimuli by either transient or persistent activation, lasting long past the input cessation. The balance between protein synthesis and dilution is crucial for establishing the time it takes to reach the critical concentrations of the regulatory proteins enabling the QS response, and thus for controlling the temporal thresholding of inputs and the timing of response initiation. This sensitivity of the initial response dynamics to the temporal fluctuations of the extracellular milieu can supply the information on environmental changes to the downstream, adaptive circuits far in advance of reaching the steady state of the QS response, which may provide additional time for eliciting appropriate responses.

Similar considerations pertain to the cases of QS involvement in symbiotic settings, as for *V. fischeri*, where the circadian cycles of animal foraging are synchronized with cycles of growth and QS response by the symbiont bacterial cells. We find that amino acid-based nutrient sources, thought to be the means of the host’s support of *V. fischeri* growth *in vivo*, may support QS response only in a certain range of temperatures, likely present in the context of ocean animals, and with highly adjustable onset time, allowing the response to be regulated by nutrients not only through control of cell density, but also more directly.

Our results also reveal a new aspect of the possible meaning of the messages exchanged by cells during QS-related cell-communication. The persistent responses of cell sub-populations following transient changes in the cell environment can serve as a type of memory of the recent exposure to conditions promoting QS, even if the environment has since become benign. This memory is carried not by the population as a whole but by different fractions of cells within the population, which can be quite limited. As mentioned above, this strategy of diversification of QS responses within a population may resemble the often discussed bet-hedging response, in which a sub-population of cells may be more adapted to a possible change in the environment than the rest of the population. However, in the case of the QS response, the non-autonomous nature imparted by cell-cell coupling through secreted AI can allow the induced subpopulation, which can anticipate the environmental change, to accelerate response by the uninduced cells, if the conditions indeed change for the worse. This allows the uninduced cells to avoid the metabolic costs associated with the initial QS induction but still enable faster onset of QS-associated adaptive response when needed. In a sense, this represents a type of ‘education’ of uninduced cells by the induced ones, providing the intercellular communication messages with a specific meaning. The resulting higher overall population responsiveness can increase the population fitness vs. the populations devoid of such cell ‘education’ capabilities.

## Methods

### Design and fabrication of microfluidic device

The overall organization of the channels and chambers in the microfluidic device is similar to that in previous studies^44,45^. Briefly, an array of 16 parallel flow-through channels with a depth of 15 μm are connected to 24 rows of 30 chambers each, which house the cells. The dimensions of the chambers are 100 × 100 × 0.8 μm, with a 15 × 15 μm post in the center. The degree to which the chambers are connected to the flow-through channels via coupling channels are varied. 16 rows of chambers are connected to flow-through channels by coupling channels only on one single side of the chamber, while eight rows of chambers are connected to flow-through channels by coupling channels on two, opposing sides of the chamber. Each side of the chamber that is connected to the flow-through channels has either one, two, or three coupling channels, giving one-sided chambers a total of one, two, or three coupling channels, while double-sided chambers have a total of two, four, or six coupling channels. In addition, the coupling channel dimensions are 25 or 50 μm in length, and 10 or 20 μm in width. The combination of different dimensions and configurations of coupling channels produced 24 unique chamber types with varying degrees of mass transport properties. The chambers are organized into four quadrants of six types each, with three single-sided and three double-sided chambers in each quadrant. Each group is then distributed in a tandem, repeated triplet fashion, to ensure that each chamber type is distributed evenly throughout the quadrant to minimize the effect of positional dependence relative to the source of medium.

Growth medium is supplied from either one of two inlets, which connect to the flow-through channels and supply nutrients to chambers on its way to a single outlet port, while the fluid from the alternate inlet is directed to a waste port. Because the height of the chambers are relatively shallow compared to the height of the flow-through channels, the chambers are much more resistant to flow across the chamber as compared to flow through the channels, hence, the dominant mode of mass transport into the chamber is through diffusion. The symmetric binary branching of the flow-through channels ensures that the pressures are balanced between channels, further preventing crossflow into the chamber. Flow is driven through the device via hydrostatic pressure differences, with the height of the syringe connected to the inlet with the desired medium higher than the alternate inlet, which are both higher than the syringes connected to the outlet and waste port. The medium being supplied to the chambers from one inlet can be instantaneously switched to that of the alternate inlet by swapping the height of the syringes, allowing for rapid changes in the medium conditions.

A chemical gradient-generating microfluidic device was also used in this study, based off of the Christmas tree design^46^. Similar to the device mentioned above, the gradient device contains 2 inlets, 1 outlet, and 1 waste port, allowing for similar operation under normal conditions. However, between the chambers and the inlets are a series of serpentine channels, which can perform mixing of solutions if medium from both inlets are supplied simultaneously. If one inlet (source) contains a molecule which the other inlet (sink) lacks, each stage of serpentine channels performs progressively more refined dilutions, resulting an increasingly resolved concentration gradient spanning the two concentrations of the inlets. Individual concentration doses are separated into channels, allowing downstream chambers to be exposed to different concentrations. The device creates 8 linearly graded concentration doses from 2 input concentrations, each connected to a row of 30 (only double-sided) or 90 (single- and double-sided) chambers, depending on the design. The design with 30 chambers in each row has only a chamber type with the highest degree of connectivity, while the design with 90 chambers in each row has 12 chamber types, all of which are contain 25 μm long coupling channels. The heights of the chamber and channels are the same as the device above, except the design with 90 chambers in each row have chamber heights of ∼0.75 μm. The channels converge into the outlet via the same symmetric binary branching structure, allowing for equal pressures. When a gradient is desired, it can be generated by equating the pressure from both syringes and supplying medium from both inlets at the same time. A fluorescent dye is used to visualize the gradient.

The device was fabricated in a similar fashion as in previous studies^44,45^. Briefly, the initial design was drawn in Adobe Illustrator and sent to be printed as a photomask. The photomask was used to produce a master mold via photolithography. The master mold was made with a 3” silicon wafer with a two-level micro-relief (0.8 μm and 15 μm) of a UV-curable epoxy (SU-8 by MicroChem, Newton, MA). The first level was made with SU-8 2002 spun onto the silicon wafer at an initial 500 rpm for 10 seconds with an acceleration index of 1, then 10000 rpm for 30 seconds with an acceleration index of 100, to produce a ∼0.8 μm thick film. The chambers were patterned with photolithography, followed by subsequent baking and development to form the structures. The subsequent level was made with SU-8 2015 spun onto the silicon wafer at an initial 500 rpm for 10 seconds with an acceleration index of 1, then 3250 rpm for 30 seconds with an acceleration index of 2 to produce a ∼15 μm thick film. The channels were patterned and made as above. Microfluidic devices were then fabricated with PDMS (Sylgard 184, Dow Corning) via soft lithography. A 5 mm thick cast of PDMS made with 10:1 ratio of elastomer base to elastomer curing agent. The PDMS cast was peeled off the wafer and cut into individual chips, and ports were punched with a 20 gauge luer stub. Devices were washed and hermetically sealed to #1.5 microscope cover slips, then baked in a 130°C oven overnight prior to use.

### Strain and growth conditions

MG1655 *E. coli* expressing a truncated Lux quorum sensing operon from *V. fischeri* made in a previous study^30^ was used. Briefly, MG01S, containing *luxR* divergently transcribed from P*luxI* fused to *GFP*, and MG02S, which is identical to MG01S bar the addition of *LuxI* upstream of *GFP* and under the same promoter, were made by the restriction digests of *Eco*RI-*Kpn*I and *Eco*RI-*Bam*HI fragments of the lux operon from pLVA01 and pLVA02, respectively, and cloned into pPROBE’-GFP-Tagless. An *Eco*RI-*Not*I digestion of the resulting plasmid and subsequent cloning into λInCh vector allowed for the genomic integration of the lux quorum sensing circuit into the *E. coli* MG1655 chromosome. 100 ug/mL ampicillin was added for selection.

The inducible LuxR strain was made by PCR amplification of the *LuxR* gene from MG02S with forward primer 5’ – ATCTCTGAATTCCCGTTTTAATGATATATAACACGCAAAACTTGCGAC – 3’, which adds an *Eco*RI restriction site upstream of the *LuxR*, and reverse primer 5’ – CAAGTATGGTACCCGTACTTAACTTTTAAAGTATGGGCAATCAATTGCTCC – 3’, which adds a *Kpn*I restriction site downstream of the gene. The PCR fragment was cloned into pEXT22 plasmid by digesting both with *Eco*RI and *Kpn*I, placing *LuxR* under the control of a tac promoter. The subsequent plasmid was transformed into MG02S. LuxR overexpression was induced with 1 mM IPTG added to growth medium. 50 ug/mL kanamycin was added for selection.

Prior to every experiment, a single colony was selected from a plate and inoculated into LB with the appropriate antibiotics and grown overnight at 30°C with shaking at 230 rpm. The next day, the overnight culture was diluted 1:100 into CACM (2% casamino acids, 1x M9 salts (12.8g Na_2_HPO_4_ 7H_2_O, 3g KH_2_PO_4_, 0.5g NaCl and 1g NH_4_Cl per liter), 1 mM MgCl_2_) with 20 mM Succinate, 20 mM Glucose and 100 ug/mL ampicillin, and grown at 30°C with shaking at 230 RPM until the culture reached an OD_600_ of 0.2-0.3 (approximately 3-4 hours). The culture was subsequently centrifuged and cell pellet re-suspended in 1% BSA in PBS prior to loading into the device. Cell loading was performed from the outlet port of a BSA in PBS-primed device. After loading into the device, growth medium with the appropriate antibiotics and 20 mM glucose is supplied to the cells continuously via one or two inlets and the cells are grown at 30°C until all chambers are full before beginning any experiment, unless otherwise indicated. Other growth media used were all variations of the CACM, with 20 mM, 10 mM, or absence of glucose, or the substitution of 2% casamino acids with 2% tryptone, both of which are sourced from digestions of casein.

### Microscopy

Widefield image acquisition was performed on a Nikon Eclipse TE2000 epifluorescence inverted microscope equipped with a motorized stage (Prior Scientific, Cambridge, UK) and a Cascade 1K EMCCD camera (Photometrics, Tucson, Arizona). Timelapse images were acquired with a Nikon Plan Fluor 40x/0.75 Ph2 DLL objective, while montage images were captured with a Nikon Plan Fluor 10x/0.3 Ph1 DL objective, and stitched together automatically with 10% image overlap with a custom MATLAB script. Both phase contrast and fluorescence images were captured for timelapse and montage images. Timelapse images were acquired every 20 minutes, with an exposure time of 100-500 ms, while the montage images were taken with an exposure time of 1000ms. The excitation filter wavelengths used for GFP detection was 450-490nm (Chroma, Rockingham, VT), while the emission filter wavelengths was 525nm. The excitation filter wavelengths used for detection of Alexafluor 555 was 541 – 565 nm, while the emission filter wavelengths was 584 – 679 nm. A spectral 2D template autofocus algorithm was used to maintain the focus for the entire duration of timelapse imaging. For time-lapsed images, at least two positions for each condition/chamber type was chosen for each experiment, while montages varied from 42 to 77 images depending on the device used. Control of the microscope and capture of images were performed in an automated fashion via Slidebook 5.5 (Intelligent Imaging Innovations, Denver, Colorado).

Z-stacks were acquired with a Leica SP8 Laser Scanning Confocal microscope equipped with a white light pulsed laser with continuous wavelength and hybrid detectors. Images were acquired on a HC PL APO 40x/1.30 Oil, CS2 objective and the Leica Application Suite X software.

### Evaluation of transport and growth in microfluidic device

The transport properties of the various chamber types were evaluated with a dye, which served as a proxy for a diffusible molecule and allowed for the visualization of the diffusion process. Alexa Fluor 555 Hydrazide (Thermo Fisher Scientific, <u>Waltham, MA</u>) was chosen as the dye due to its relatively high molecular weight as compared to most diffusible molecules of interest (1150, vs 213.23g/mol for AI and 180.16g/mol for glucose), giving it a smaller diffusion coefficient and longer diffusion time. Thus, it served as a good representation of the dynamics of even the slowest diffusing molecules while also accounting for the dynamics of a wide range of molecules with faster diffusion dynamics. Cells were grown in dye-free medium within the chambers overnight until the chambers were filled to capacity. 0.01 mM of dye solution was perfused into the flow-through channels by increasing the hydrostatic pressure of the dye-containing medium syringe relative to the dye-free medium syringe. Fluorescence timelapse microscopy was used to capture the fluorescence intensity within the chambers, at a rate of 1 minute per time point, for a duration of 15 minutes. The diffusion of the dye out of the chamber was also monitored by switching back to the dye-free medium syringe. The intensity of the fluorescence in the chamber is quantified by taking the mean of the intensity of the entire chamber for each time point, and normalized to the initial or final timepoint, for diffusion out and diffusion in, respectively. The transport properties of the various chamber types was also modeled computationally with COMSOL Multiphysics 4.2 (COMSOL Inc. Stockholm, Sweden), the results of which was in agreement with experiments.

The growth rate of the cells within the different chambers was assessed with the MG01S strain, which lacks the ability to produce LuxI. Therefore, the production of high levels of GFP relies on the addition of exogenous AI, and ceases upon its removal. MG01S is loaded into chambers and grown in the presence of 1 µM AI to induce uniform GFP production until all chambers are filled to capacity. Once AI is removed, fluorescence in the chamber is gradually lost as the continual growth of the cells within the chamber would lead to the distribution of a fixed number of GFP protein to the progeny, which are gradually pushed out of the chamber by the growing and dividing cells within the chamber and carried away by the flow. Due to the stability of the GFP used^47^, the loss of GFP fluorescence is expected to be predominantly from dilution of the protein as a result of growth and division of the cells, the rate of which would be proportional to the growth rate within different chambers. The average fluorescence over time can be fitted with an exponential curve of the form a*exp(bt), giving an estimate of the growth rate. The growth rate was estimated for all 24 chamber types with both casamino acids and tryptone-based minimal media, and at 25 and 30°C.

### Image processing and data analysis

Automated image acquisition was performed with Slidebook 5.5. Raw images were exported in .tiff format and processed using custom MATLAB scripts. All images were corrected for uneven illumination with the following correction: C = (I-D)/(F-D)*M, where C is the corrected image, I is the initial image, D is the darkfield image, F is the flatfield image, and M is the mean of the flatfield minus darkfield images. The flatfield and darkfield were averages of multiple images. Time-lapse images were aligned automatically with an alignment algorithm, while montage images were stitched together with the same alignment algorithm with 10% overlap between adjacent images. Each calculated metric is an average of at least 3 independent experiments of at least 2 replicates each for each chamber configuration, unless otherwise indicated.

The automated data analysis pipeline imports the processed image file and determines the square chamber region, excluding the post, containing cells. The mean intensity value for the entire chamber was determined for both sets of image types. For time-lapse images, this is done for all time points, and additional metrics were also calculated. To visualize the spatial distribution of response along the y-axis, the intensity values were averaged along the x-axis and the resultant column of intensity values were normalized by subtracting the minimum and dividing by the minimum-subtracted maximum. The steepness of spatial gradients at each time point was approximated with the slope from a linear regression performed on the column of intensity values versus a normalized distance of 1, with a positive slope representative of higher responses in the distal regions relative to the coupling channels, and a negative slope representative of higher responses in the proximal regions relative to the coupling channels. Double-sided chambers are analyzed as half-chambers along the y-axis due to symmetry, and the calculated metric for both halves are averaged into a single value to represent the whole chamber. Aggregating these intensity value columns for each time point generated the kymographs used for spatial analysis. Onset time is defined as the time point at which more than 1.5% of the pixels within the square region of the chamber have intensity values greater than mean + 4*SD of the same region in the initial frame. Onset location was quantified from the kymographs. Briefly, the columns of normalized intensity values from the onset time point to a time point 2 hours later were extracted, and the positions with a normalized value of greater than or equal to 0.99 were collected and the mean and standard deviations were calculated. The initial rate was estimated with the slope from a linear regression on the first 5 hours of response after the onset time within exogenous induction conditions, which is expected to reach maximal levels of production. Time-lapse fluorescence intensity measurements are normalized to the initial value. Two chamber configurations were excluded (filled shapes) from the chip-wide measurements obtained from the montage image analysis due to partial formation of cell bilayer, but z-sections from confocal microscopy confirm the biphasic nature (Fig. S3). Fractions of induction within a chamber were determined by the fraction of total pixels that exceeded a predetermined threshold value. For categorization, mean±SD ≤10% and ≥0.01% were considered partially induced, less than that is uninduced, and greater than that is fully induced. For chambers that are not packed at the beginning of the experiment, segmentation of cells based on phase contrast images were performed, and all quantified data from these chambers were from the segmented images.

Z-stacks from confocal microscopy were split into individual focal planes, with the chamber region for each plane segmented and the mean intensity calculated. The slice with the highest mean intensity was used for analysis, and were compared to the mean value from the maximum intensity projections.

### Computation of mutual information

Briefly, mutual information (*I*) between two random variables is calculated from the formula:

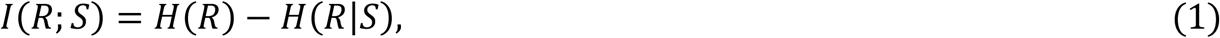

where *S* and *R* denote the signal and response, respectively, and *H* is the entropy function:

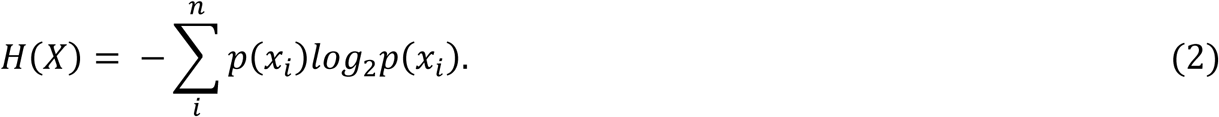

Hence, formula (1) becomes:

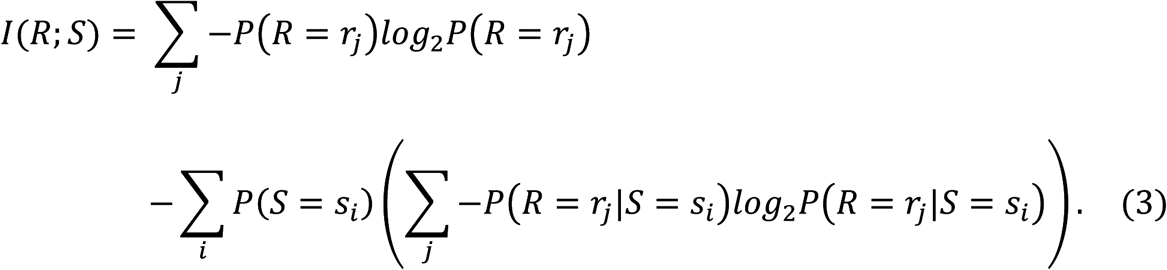

The marginal distribution of the response is

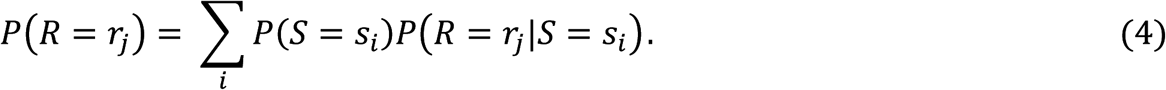

Since the formulas as shown above are for discrete data, the experimental data used for the calculations, which lies on a continuous spectrum, are discretized by binning. However, binning of the finite data sample results in biased estimates of the mutual information. Since bias is a function of sample size, which approaches zero as sample size approaches infinity, we utilized the series expansion of mutual information in terms of inverse sample size to estimate the unbiased mutual information:

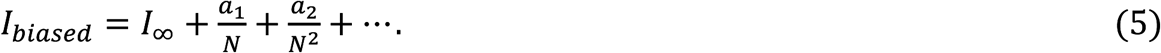

Here, *I*_*baised*_ is the biased estimate of mutual information, *I*_∞_ is the unbiased estimate of mutual information with infinite sample size, N is the total number of samples, and *a*_*i*_ are coefficients dependent on the signal and response distributions. For sufficiently large N, all terms larger than first order are negligible, resulting in the biased estimate of mutual information being a linear function of inverse sample size. As a result, we can use jackknife sampling to sample subsets of the data to compute the mutual information. By plotting these biased estimates of mutual information to the inverse sample size, fitting a line, and extrapolating to infinite sample size, we can obtain the unbiased estimate of mutual information. For a more comprehensive description, we refer the reader to previously published work^35,36^.

### Model and simulations

#### Simulation of diffusion

Diffusion dynamics were simulated using COMSOL Multiphysics 4.2. Briefly, the 3D geometry of each chamber configuration was recreated within the software, and the concentration of the entire volume of the chamber was set to an initial concentration of 0 mM. All surfaces of the chamber were set to have null flux except for the surfaces that interface with the flow-through channels, which are set to a constant concentration of 0.01 mM, corresponding to the concentration of dye used in the experiment. A diffusion coefficient of 2 × 10^−10^ *m*^2^ *s*^−1^ was used for the Alexa Fluor 555 dye. Time-dependent transport of diluted species simulations were performed for all 24 chamber configurations. The average concentration within the chamber volume was determined and plotted.

#### Simulation of QS response

The mathematical model is a simplified version of a model from a previous study^34^, accounting for only one cell, but with new species and terms added:

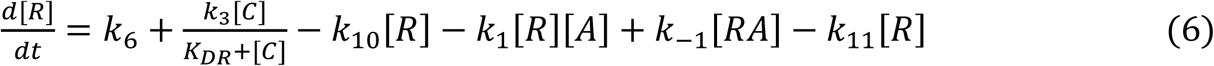

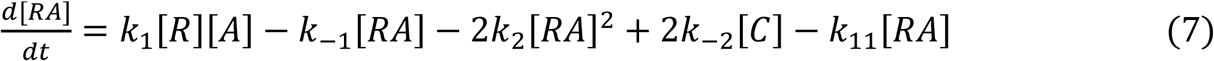

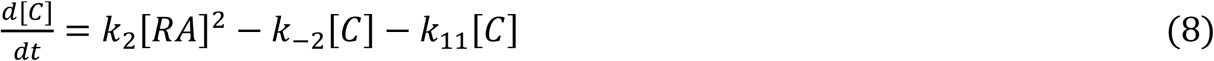

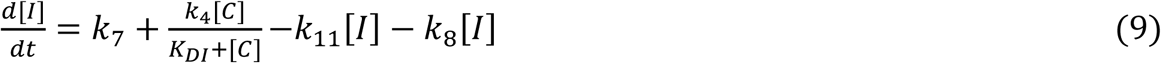

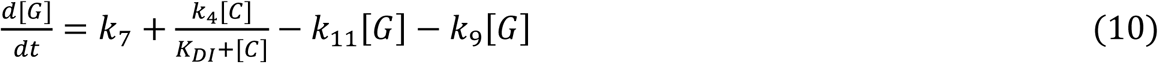

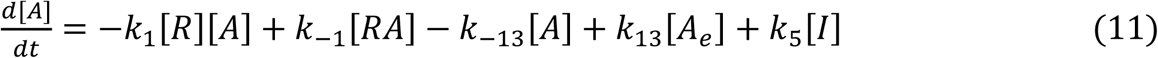

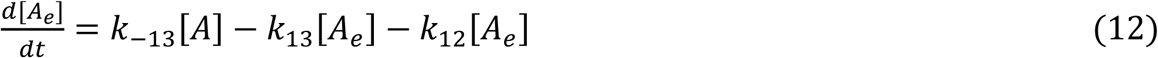

Diffusion of AI is only considered as a rate of loss from the cell, if no exogenous AI is introduced. In addition, the production of AI from C has been explicitly separated into production of LuxI from C (which then produces AI) to enable the account of LuxI dilution. Because GFP is downstream of LuxI, the added equation is nearly identical, with a different degradation term the only difference. Dilution terms were also added to the equations of all protein species to account for loss of protein from cell division.

The global constants that are applicable to all the conditions are listed in Table 1, most of which were based on a previous study^33^. For each environmental condition, a new set of k_4_ (synthesis, Table 2) and k_11_ (dilution, Table 3) values were used, the range of which corresponds to the different chamber configurations, and were measured from experiments. k_4_ (synthesis) was estimated with the slope from a linear regression corresponding to the first 5 hours of response after the onset time within exogenous induction conditions, which is expected to reach maximal levels of production. Dilution rates are proportional to growth rate, so k_11_ (dilution) is calculated in the same way with an exponential fit of the GFP dilution data. The exceptions are k_4_ (synthesis) for tryptone and casamino acids conditions at 30°C, which were inferred by comparing the change in growth rates and synthesis rates from tryptone to casamino acids medium at 25°C, and extrapolating that relationship to the change in temperature from 25°C to 30°C in both media types.

Simulations were performed in MATLAB. Briefly, each simulation was run for an initial 24 simulation hours with either the LuxR repressed or LuxR partially repressed k_3_ and k_6_ values, to replicate the initial growth in 20 mM or 10 mM glucose medium, respectively, and to allow for a basal steady state amount of proteins and components to be made. The k_3_ and k_6_ values are subsequently switched to the derepressed or overexpressed form, and simulation resumed for at least another 24 simulation hours. Simulations corresponding to the transient glucose switching experiments consisted of transient changes in the k_3_ and k_6_ values at the appropriate time points.

## Statistical Analysis

At least 3 independent tests were performed for each experiment unless otherwise stated. F-test was performed to determine variances between samples. 2-tailed t-tests were used for comparisons.

## Code Availability

The codes used in this study are available from the corresponding author upon request.

## Data Availability

The data that support the findings of this study are available from the corresponding author upon request.

## Acknowledgements

We thank A. Stevens for the generous gifts of the cell strain used in this study. We are grateful to everyone in the Levchenko group for the helpful discussions.

## Author Contributions

E.K.C., A.G. and A.L. designed the device and E.K.C. fabricated the device. E.K.C. performed experiments and simulations, and analyzed data. E.K.C. and A.L. wrote the paper.

