## Supplementary info for "Environmental sensing in dynamic quorum responses"

### Table 1: Model parameters

| Symbol | Value | Units | Description | Reference |
|---|---|---|---|---|
| $k_1$ | $5 \times 10^{-5}$ | $m^{-1} s^{-1}$ | Association between R and A | (1) |
| $k_{-1}$ | $5 \times 10^{-4}$ | $s^{-1}$ | Dissociation of RA | (1) |
| $k_2$ | $1 \times 10^{-5}$ | $m^{-1} s^{-1}$ | Dimerization of RA | (1) |
| $k_{-2}$ | 0.01 | $s^{-1}$ | Dissociation of C | (1) |
| $k_3$ (LuxR repressed, +20 mM Glc) | $7 \times 10^{-4}$ | $m s^{-1}$ | Feedback production of R, maximal rate | Estimated |
| $k_3$ (LuxR partially repressed, +10 mM Glc) | 0.00535 | $m s^{-1}$ | Feedback production of R, maximal rate | Estimated |
| $k_3$ (LuxR derepressed, -Glc) | 0.01 | $m s^{-1}$ | Feedback production of R, maximal rate | Estimated |
| $k_3$ (LuxR Overexpressed) | 0.1 | $m s^{-1}$ | Feedback production of R, maximal rate | Estimated |
| $k_4$ | Variable, See Table 2 | $m s^{-1}$ | Feedback production of I, maximal rate | Measured |
| $k_5$ | 0.45 | $s^{-1}$ | Enzymatic production of A | (1) |
| $k_6$ (LuxR repressed, +20 mM Glc) | $7 \times 10^{-4}$ | $m s^{-1}$ | Constitutive production of R | (1) |
| $k_6$ (LuxR partially repressed, +10 mM Glc) | 0.00285 | $m s^{-1}$ | Constitutive production of R | (1) |
| $k_6$ (LuxR derepressed, -Glc) | 0.005 | $m s^{-1}$ | Constitutive production of R | (1) |
| $k_6$ (LuxR Overexpressed) | 0.05 | $m s^{-1}$ | Constitutive production of R | Estimated |
| $k_7$ | $1 \times 10^{-5}$ | $m s^{-1}$ | Constitutive production of I | Estimated |
| $k_8$ | $5 \times 10^{-5}$ | $s^{-1}$ | Degradation of I | (1) |
| $k_9$ | $7.4 \times 10^{-6}$ | $s^{-1}$ | Degradation of GFP | (2) |
| $k_{10}$ | 0.001 | $s^{-1}$ | Degradation of R | (1) |
| $k_{11}$ | Variable, See Table 3 | $s^{-1}$ | Dilution rate | Measured |
| $k_{12}$ | 0.01 | $s^{-1}$ | Removal of extracellular A | Estimated |
| $k_{13}$ | 0.4 | $s^{-1}$ | Exchange of A from extracellular to intracellular | (1) |
| $k_{-13}$ | 0.4 | $s^{-1}$ | Exchange of A from intracellular to extracellular | (1) |
| KDI | 25 | m | Dissociation constant for I | (1) |
| KDR | 1 | m | Dissociation constant for R | (1) |

### Table 2: Dilution rates (Order defined based on Tryptone 25°C condition) ($s^{-1}$)

| | Tryptone 25°C | Casamino Acids 25°C | Tryptone 30°C | Casamino Acids 30°C |
|---|---|---|---|---|
| singlesinglelongnarrow | 9.02E-06 | 3.63E-07 | 1.05E-05 | 5.43E-06 |
| doublesinglelongnarrow | 1.17E-05 | 1.91E-06 | 1.14E-05 | 6.99E-06 |
| triplesinglelongnarrow | 1.24E-05 | 4.68E-06 | 1.26E-05 | 1.11E-05 |
| singledoublelongnarrow | 1.37E-05 | 6.02E-06 | 1.34E-05 | 1.29E-05 |
| singlesingleshortnarrow | 1.40E-05 | 7.19E-06 | 2.18E-05 | 1.92E-05 |
| singlesinglelongwide | 1.50E-05 | 8.49E-06 | 2.36E-05 | 2.18E-05 |
| doubledoublelongnarrow | 1.90E-05 | 1.01E-05 | 2.1E-05 | 1.82E-05 |
| doublesingleshortnarrow | 2.14E-05 | 1.38E-05 | 2.95E-05 | 2.71E-05 |
| singlesingleshortwide | 2.30E-05 | 1.73E-05 | 3.46E-05 | 3.51E-05 |
| doublesinglelongwide | 2.57E-05 | 1.80E-05 | 3.43E-05 | 3.54E-05 |
| triplesingleshortnarrow | 2.84E-05 | 1.83E-05 | 3.48E-05 | 3.77E-05 |
| tripledoublelongnarrow | 2.89E-05 | 2.05E-05 | 3.02E-05 | 2.91E-05 |

| | | | | |
|---|---|---|---|---|
| triplesinglelongwide | 3.03E-05 | 2.03E-05 | 4.05E-05 | 4.07E-05 |
| singledoublelongwide | 3.42E-05 | 2.36E-05 | 4.87E-05 | 4.75E-05 |
| singledoubleshortnarrow | 3.50E-05 | 2.29E-05 | 5.16E-05 | 4.33E-05 |
| doublesingleshortwide | 3.63E-05 | 2.59E-05 | 4.94E-05 | 5.59E-05 |
| triplesingleshortwide | 3.95E-05 | 2.80E-05 | 5.65E-05 | 5.83E-05 |
| doubledoubleshortnarrow | 4.57E-05 | 3.03E-05 | 6.44E-05 | 6.08E-05 |
| doubledoublelongwide | 4.96E-05 | 3.31E-05 | 6.62E-05 | 6.92E-05 |
| singledoubleshortwide | 4.99E-05 | 3.42E-05 | 7.32E-05 | 7.35E-05 |
| tripledoubleshortnarrow | 5.31E-05 | 3.54E-05 | 7.72E-05 | 7.13E-05 |
| tripledoublelongwide | 5.39E-05 | 3.65E-05 | 7.9E-05 | 7.88E-05 |
| doubledoubleshortwide | 5.92E-05 | 4.04E-05 | 9.29E-05 | 8.43E-05 |
| tripledoubleshortwide | 6.22E-05 | 4.36E-05 | 0.000104 | 8.56E-05 |

**Table 3: Synthesis rate (Order defined based on Tryptone 25°C condition) (m $s^{-1}$)**

| | Tryptone 25°C | Casamino Acids 25°C | Tryptone 30°C (extrapolated) | Casamino Acids 30°C (extrapolated) |
|---|---|---|---|---|
| singlesinglelongnarrow | 0.006953 | 0.016613 | 0.001 | 0.014001 |
| doublesinglelongnarrow | 0.010404 | 0.0273 | 0.00169 | 0.014543 |
| triplesinglelongnarrow | 0.015151 | 0.032305 | 0.00238 | 0.015085 |
| singledoublelongnarrow | 0.01542 | 0.031761 | 0.00307 | 0.015627 |
| singlesingleshortnarrow | 0.015919 | 0.025956 | 0.003761 | 0.016169 |
| singlesinglelongwide | 0.016091 | 0.025045 | 0.004451 | 0.016711 |
| doubledoublelongnarrow | 0.022878 | 0.037048 | 0.005141 | 0.017253 |
| doublesingleshortnarrow | 0.017174 | 0.02727 | 0.005831 | 0.017795 |
| singlesingleshortwide | 0.019349 | 0.028035 | 0.006521 | 0.018337 |
| doublesinglelongwide | 0.018987 | 0.029261 | 0.007211 | 0.018879 |
| triplesingleshortnarrow | 0.018061 | 0.02931 | 0.007902 | 0.019421 |
| tripledoublelongnarrow | 0.025332 | 0.035974 | 0.008592 | 0.019963 |
| triplesinglelongwide | 0.022206 | 0.030584 | 0.009282 | 0.020505 |
| singledoublelongwide | 0.021332 | 0.03361 | 0.009972 | 0.021047 |
| singledoubleshortnarrow | 0.019509 | 0.034323 | 0.010662 | 0.021589 |
| doublesingleshortwide | 0.021258 | 0.034675 | 0.011352 | 0.022131 |
| triplesingleshortwide | 0.024263 | 0.036005 | 0.012043 | 0.022674 |
| doubledoubleshortnarrow | 0.026939 | 0.041492 | 0.012733 | 0.023216 |
| doubledoublelongwide | 0.032675 | 0.042832 | 0.013423 | 0.023758 |
| singledoubleshortwide | 0.031864 | 0.041858 | 0.014113 | 0.0243 |
| tripledoubleshortnarrow | 0.038133 | 0.045328 | 0.014803 | 0.024842 |
| tripledoublelongwide | 0.039596 | 0.046597 | 0.015493 | 0.025384 |
| doubledoubleshortwide | 0.041151 | 0.048342 | 0.016184 | 0.025926 |
| tripledoubleshortwide | 0.038491 | 0.048087 | 0.016874 | 0.026468 |


1 Goryachev, A. B., Toh, D. J. & Lee, T. Systems analysis of a quorum sensing network: design constraints imposed by the functional requirements, network topology and kinetic constants. Biosystems 83, 178-187, doi:10.1016/j.biosystems.2005.04.006 (2006).

2 Miller, W. G., Leveau, J. H. & Lindow, S. E. Improved gfp and inaZ broad-host-range promoter-probe vectors. Mol Plant Microbe Interact 13, 1243-1250, doi:10.1094/MPMI.2000.13.11.1243 (2000).

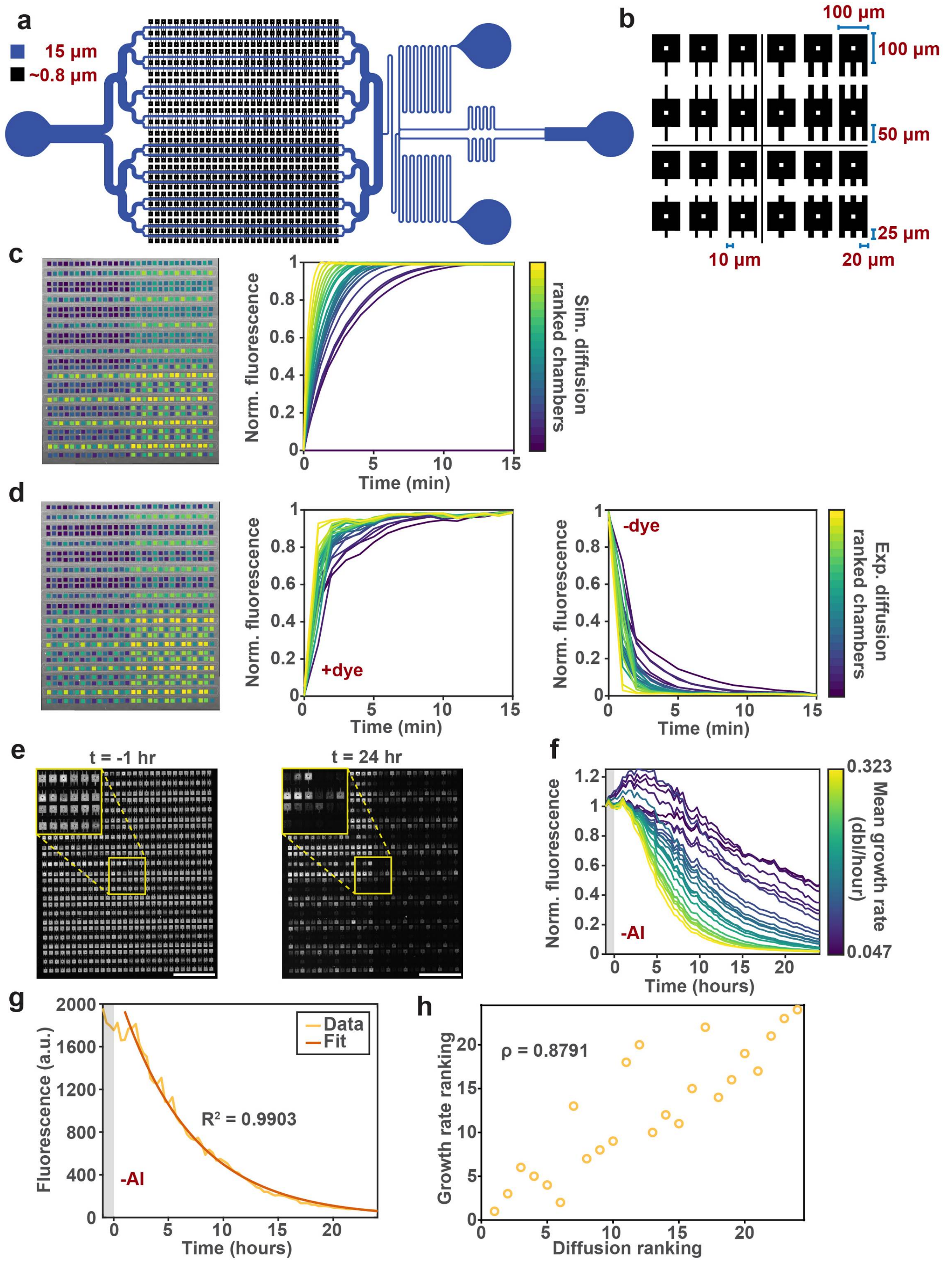
a
15 µm
~0.8 µm
b
100 µm
100 µm
50 µm
25 µm
10 µm
20 µm
c
Norm. fluorescence
Time (min)
Sim. diffusion ranked chambers
d
+dye
-dye
Norm. fluorescence
Time (min)
Exp. diffusion ranked chambers
e
t = -1 hr
t = 24 hr
f
Norm. fluorescence
-AI
Time (hours)
Mean growth rate (dbl/hour)
0.323
0.047
g
Fluorescence (a.u.)
Data
Fit
R² = 0.9903
-AI
Time (hours)
h
ρ = 0.8791
Growth rate ranking
Diffusion ranking

**Supplementary Figure 1 Characterization of microfluidic platform. a,** Diagram of microfluidic device. **b**, Diagram of the 24 unique chamber configurations, grouped into the four quadrants of the chamber array. **c,d,** Phase contrast micrograph of microfluidic device color-coded according to the rankings of **(c)** simulated and **(d)** experimental dye diffusion dynamics within chambers. Experimental dynamics for both diffusion into and out of the chamber are shown (mean, n=2). **e,** Fluorescence micrographs of chamber array containing MG01S cells taken 1 hour before and 24 hours after AI removal in tryptone medium at 25°C. Insets contain magnified view of the region indicated, which contains all 24 chamber configurations. Scale bars, 1 mm. **f,** Protein dilution dynamics in chambers used for growth rate calculation (mean, n = 6, from 3 independent experiments). **g,** Representative example of signal decay overlaid with exponential fit after AI removal. **h**, Correlation of chamber rankings determined from experimental diffusion dynamics and growth rate.

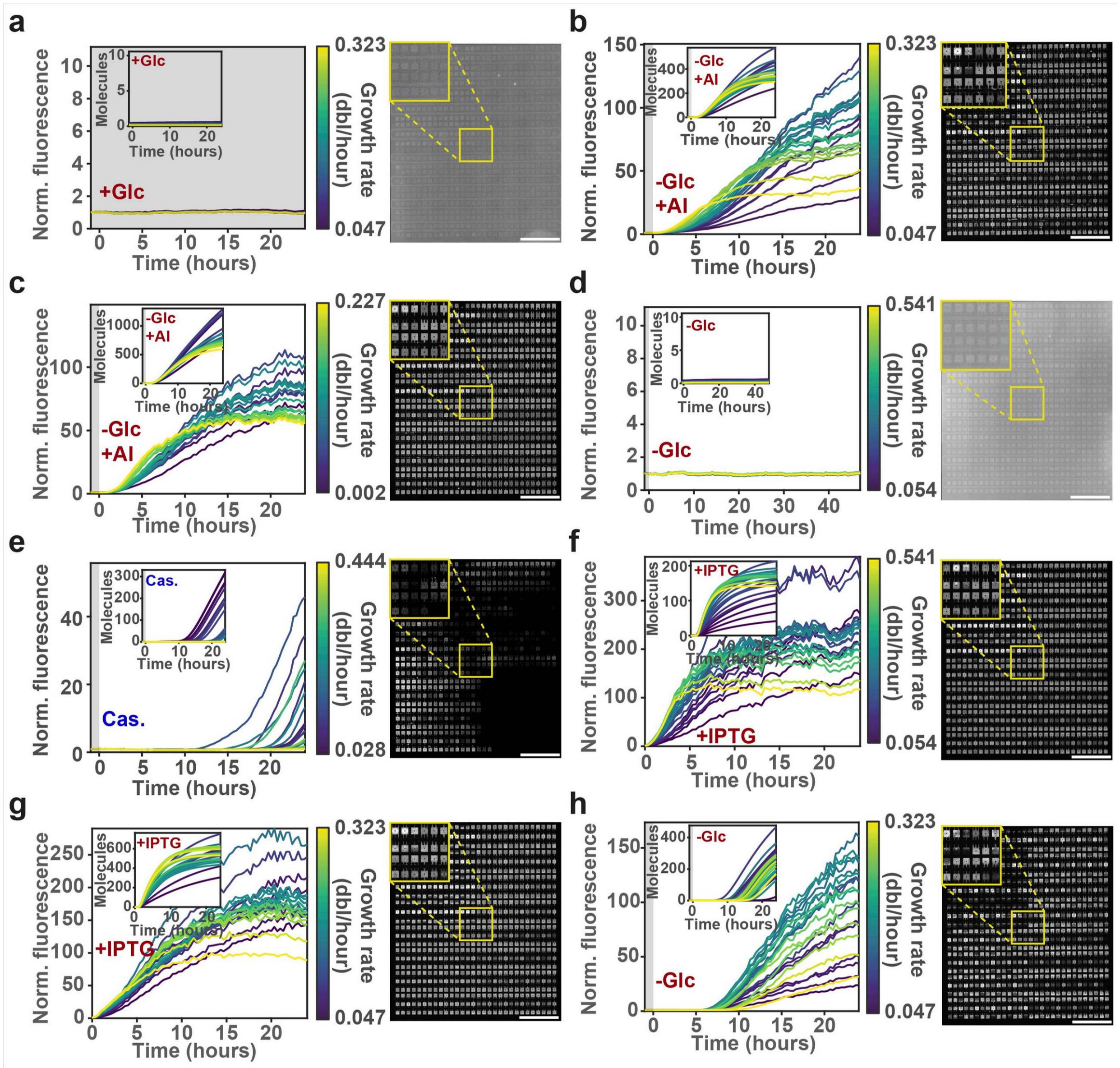


**Supplementary Figure 2 QS response within various environmental conditions. a-g,** Response dynamics and fluorescence micrograph of various conditions: **(a)** tryptone medium with 20 mM glucose at 25°C, **(b)** tryptone medium with 1 μM exogenous AI at 25°C, **(c)** casamino acids medium with 1 μM exogenous AI at 25°C, **(d)** tryptone medium at 30°C, **(e)** casamino acids medium at 30°C, **(f)** tryptone medium with 1 mM IPTG (for LuxR overexpression) at 30°C, **(g)** tryptone medium with 1 mM IPTG (for LuxR overexpression) at 25°C, and **(h)** tryptone medium with 10 mM glucose at 25°C. Inset in plot contains simulated dynamics. Inset in fluorescence micrograph contains magnified view of the region indicated, which contains all 24 chamber configurations. Mean, n = 6, from 3 independent experiments. Scale bars, 1 mm.

a

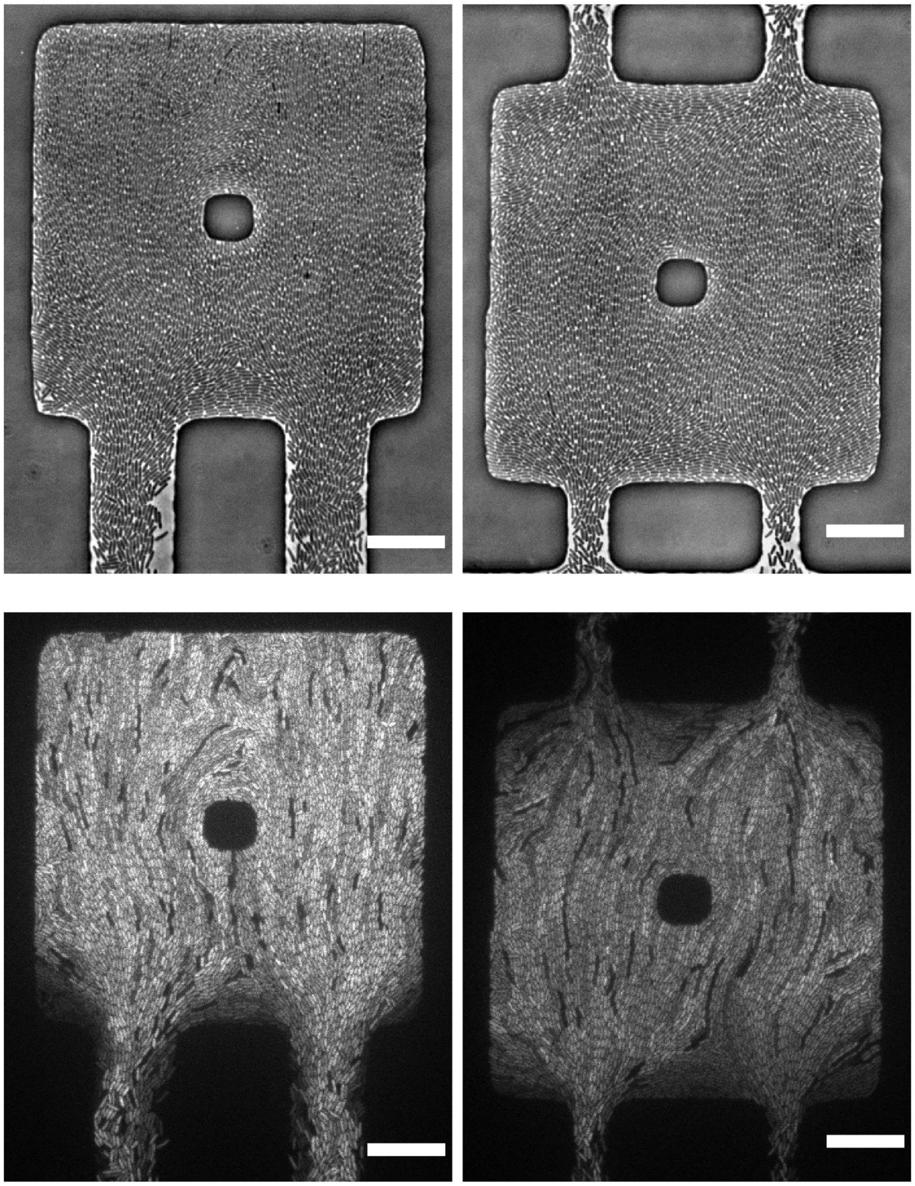

b

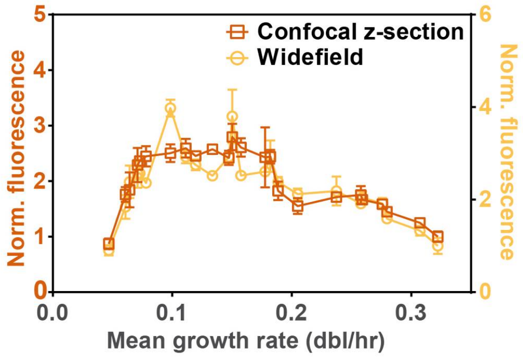


**Supplementary Figure 3 Response variation is inherent at the single-cell level. a,** Phase contrast and fluorescence micrographs of representative chambers grown in tryptone-based media at 25°C. Scale bar, 20 μm. **b,** 24$^{th}$ hour QS response distributions from tryptone medium condition with 1 μM exogenous AI at 25°C, quantified from confocal z-sections (n=2) and widefield images. (n=105 or n=60, for single- or double-sided chambers, respectively, from 3 independent experiments) (mean ± SD)

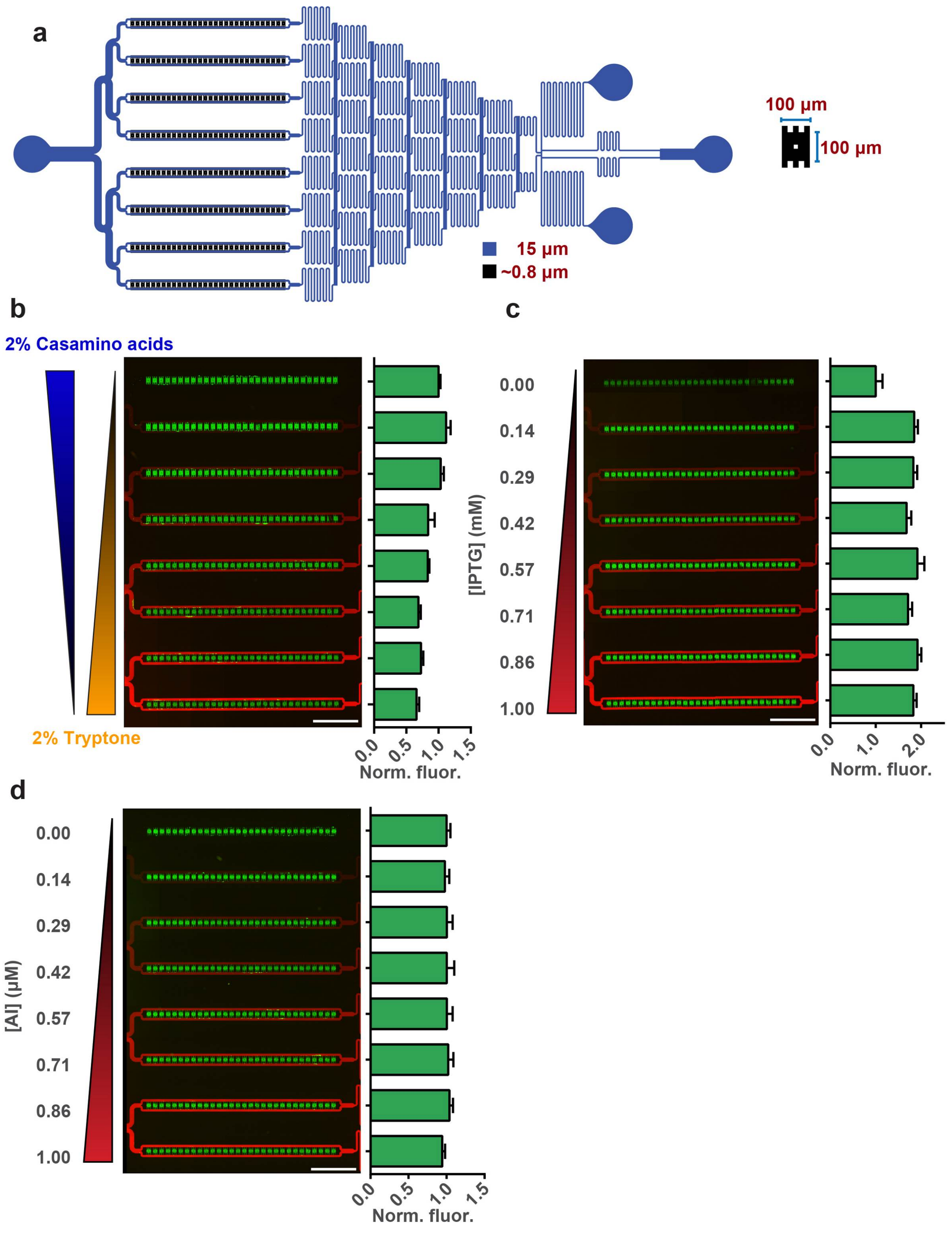


**Supplementary Figure 4 Juxtaposition of environmental conditions within gradient-generating device. a,** Diagram of gradient-generating microfluidic device with a singular chamber configuration. **b**, Fluorescence micrograph and distribution of QS response in opposing tryptone and casamino acids gradients with 1 μM exogenous AI at 25°C. **c,** Fluorescence micrograph and distribution of QS response in tryptone medium with 0-1 mM IPTG gradient (for LuxR overexpression) at 25°C. **d**, Fluorescence micrograph and distribution of QS response in tryptone medium with 0-1 μM AI gradient at 25°C. Scale bars, 1 mm. (n=30) (mean ± SD)

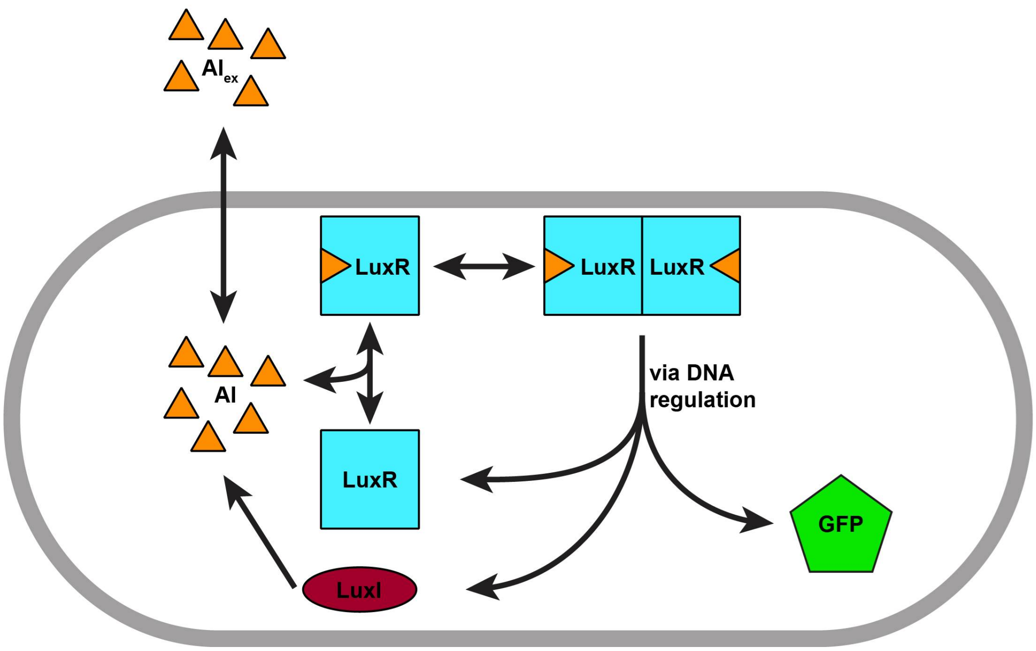


**Supplementary Figure 5 Diagram of model components**

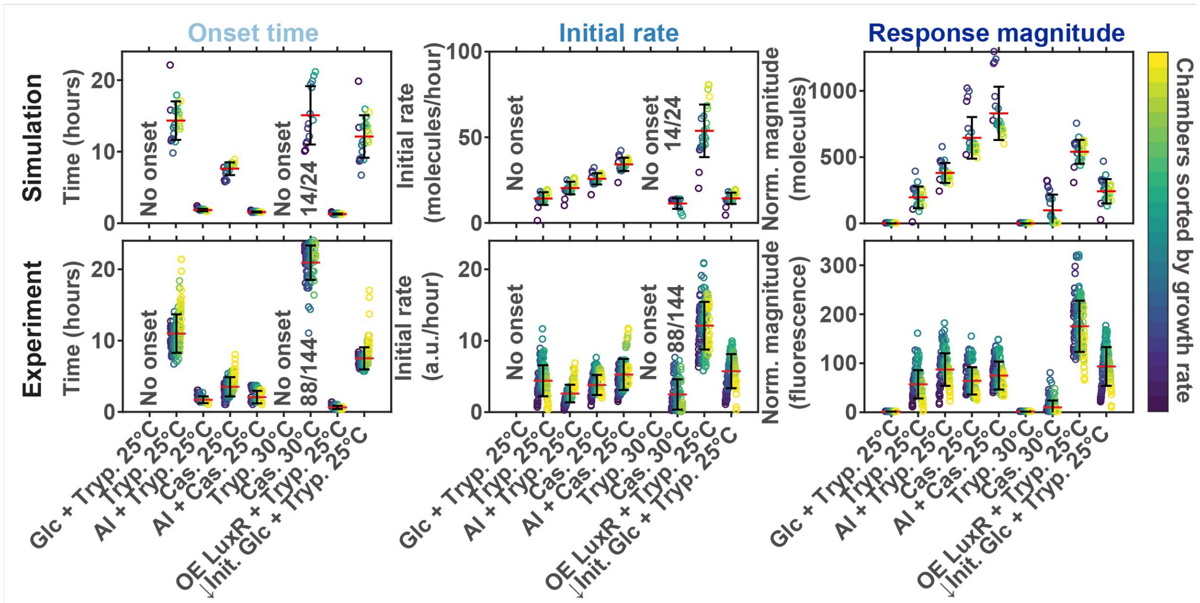


**Supplementary Figure 6 Summary of simulated and experimental QS response dynamics in various conditions.** Onset time, initial response rate, and 24$^{th}$ hour response distribution from mathematical model and experiments for all conditions tested. (n = 24 for mathematical model, n = 144 for experiment, from 3 independent experiments) (mean ± SD)

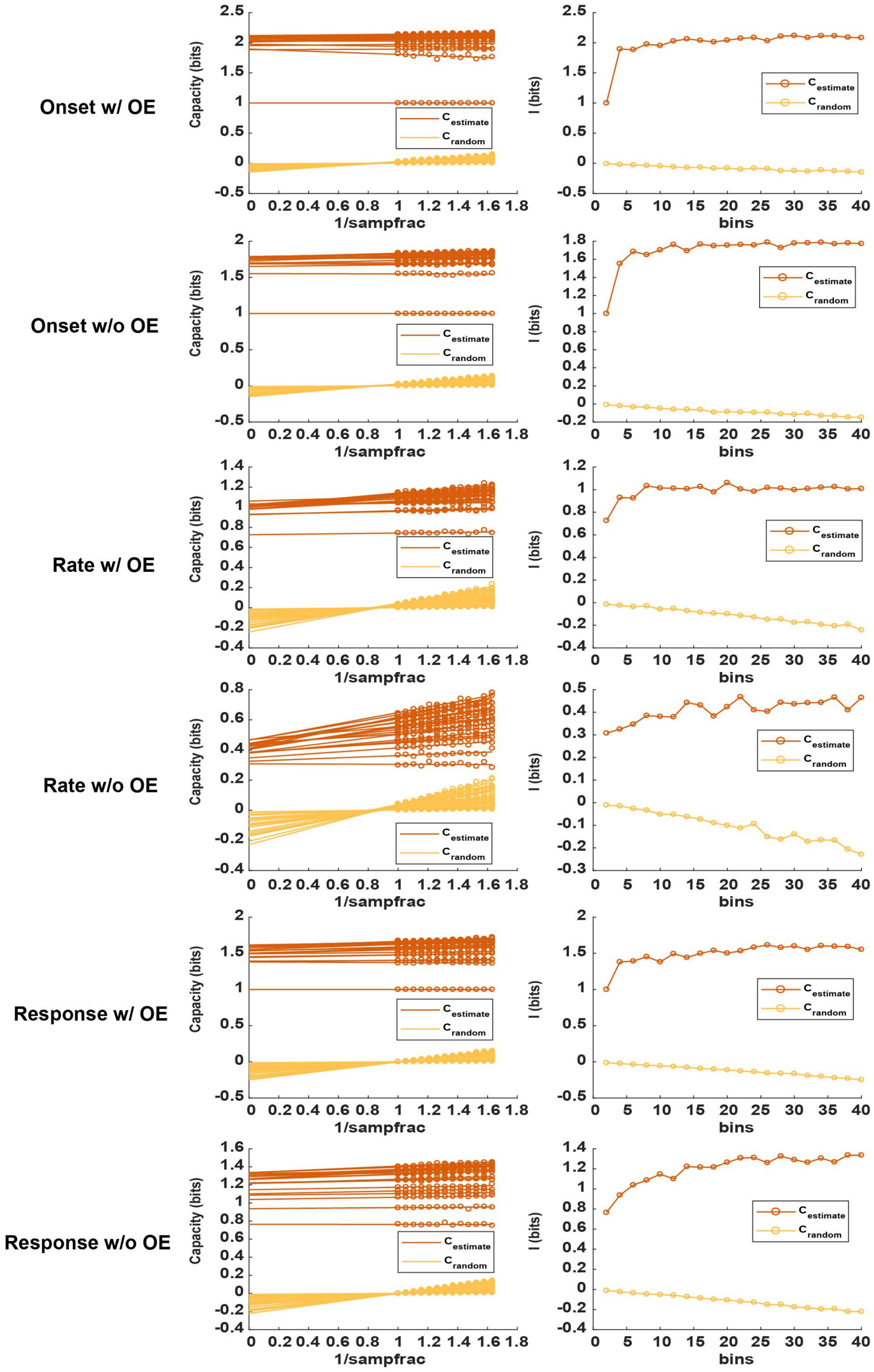

Onset w/ OE
Onset w/o OE
Rate w/ OE
Rate w/o OE
Response w/ OE
Response w/o OE
Capacity (bits)
1/sampfrac
I (bits)
bins
$C_{estimate}$
$C_{random}$

**Supplementary Figure 7 Determination of unbiased mutual information about global environmental conditions for various metrics.** Linear extrapolation to infinite sample size to determine unbiased estimate of mutual information about global environmental conditions for various metrics.

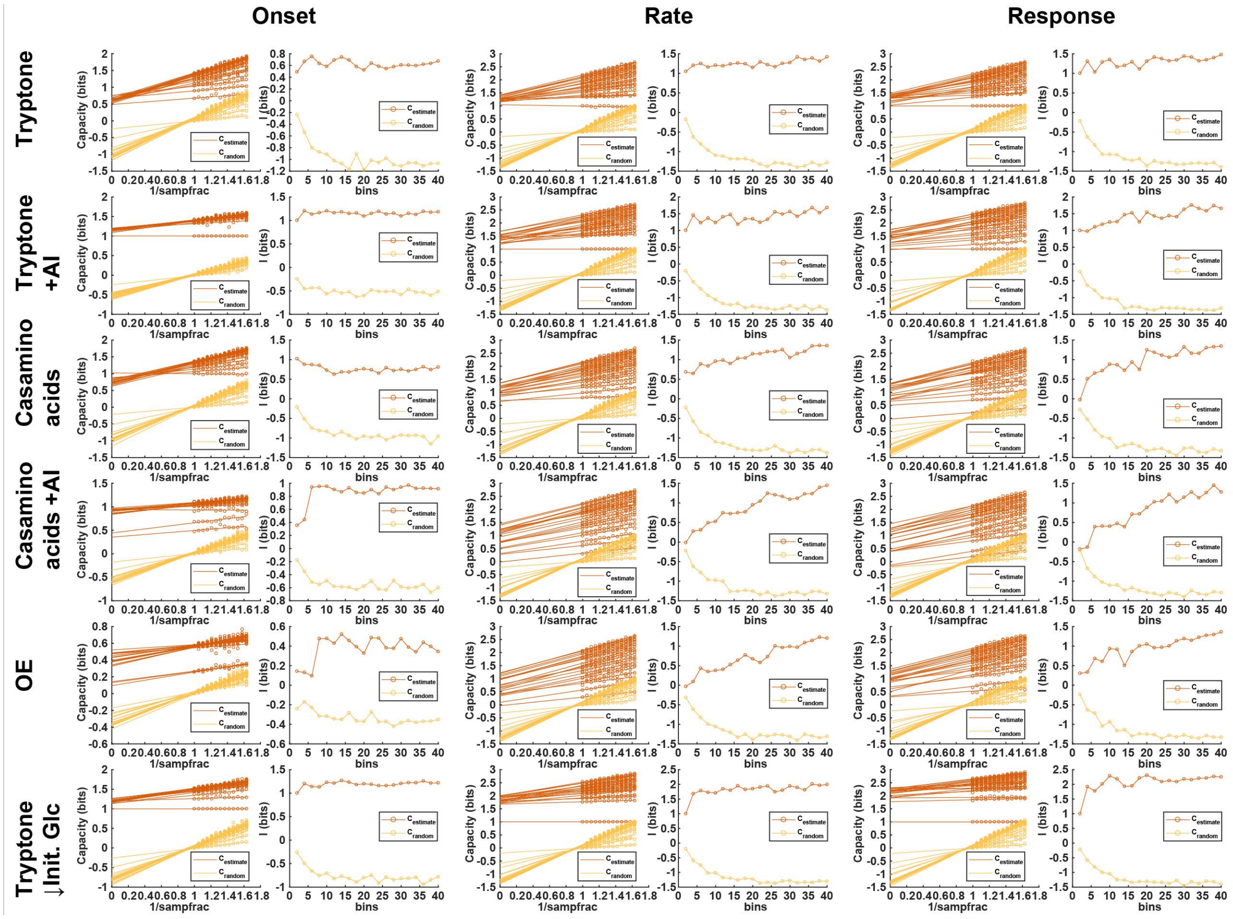


**Supplementary Figure 8 Determination of unbiased mutual information about local environmental conditions for various metrics.** Linear extrapolation to infinite sample size to determine unbiased estimate of mutual information about local environmental conditions for various metrics.

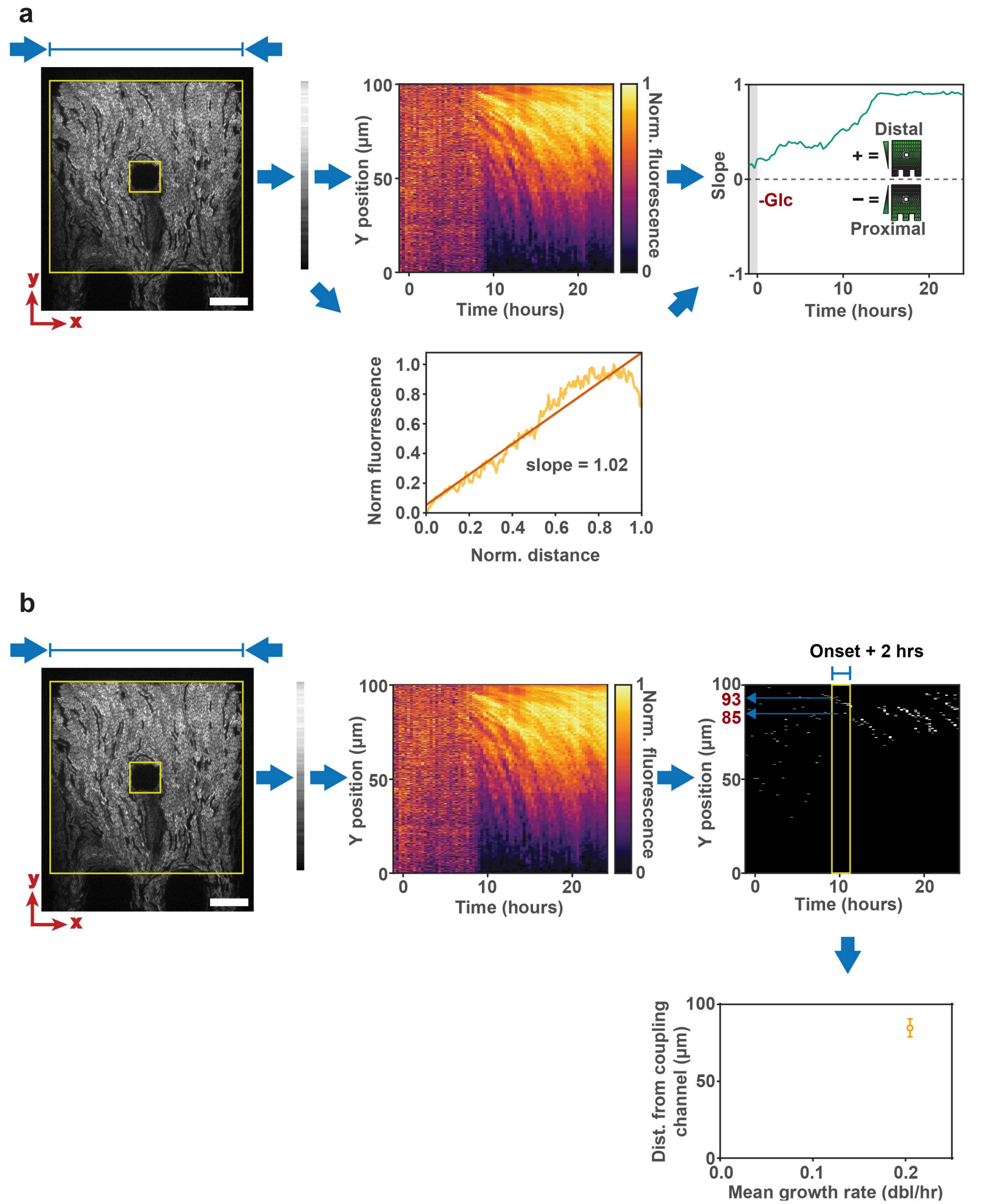


**Supplementary Figure 9 Workflow for quantification of spatial response gradient and onset location. a,b** Diagrams illustrating the workflow for the quantification of **(a)** spatial response gradient and **(b)** spatial onset localization within chambers. Scale bar, 20 μm.

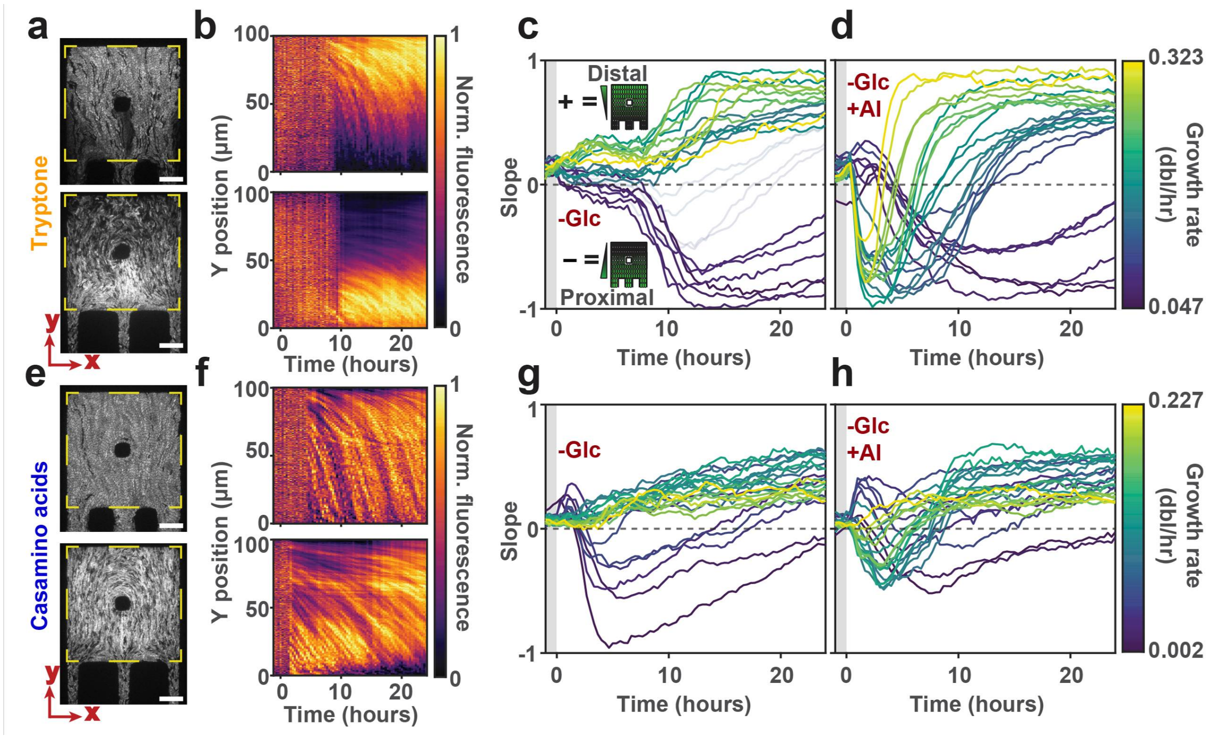


**Supplementary Figure 10 Quantification of spatial response gradients in various conditions. a,b,** Representative **(a)** fluorescence micrographs and **(b)** kymographs of chambers grown in tryptone medium at 25°C. Scale bar, 20 µm. **c,d,** Quantification of spatial response distributions with linear regression in **(c)** tryptone and **(d)** tryptone with 1 µM exogenous AI medium at 25°C. Positive slope values indicate stronger response in the regions distal from the coupling channels, whereas negative slope values indicate stronger response in the regions proximal from the coupling channels. (n = 6, from 3 independent experiments). **e,f,** Representative **(e)** fluorescence micrographs and **(f)** kymographs of chambers grown in casamino acids medium at 25°C. Scale bar, 20 µm. **g,h,** Quantification of spatial response distributions with linear regression in **(g)** casamino acids and **(h)** casamino acids with 1 µM exogenous AI medium at 25°C. Positive slope values indicate stronger response in the regions distal from the coupling channels, whereas negative slope values indicate stronger response in the regions proximal from the coupling channels. (mean, n = 6, from 3 independent experiments).

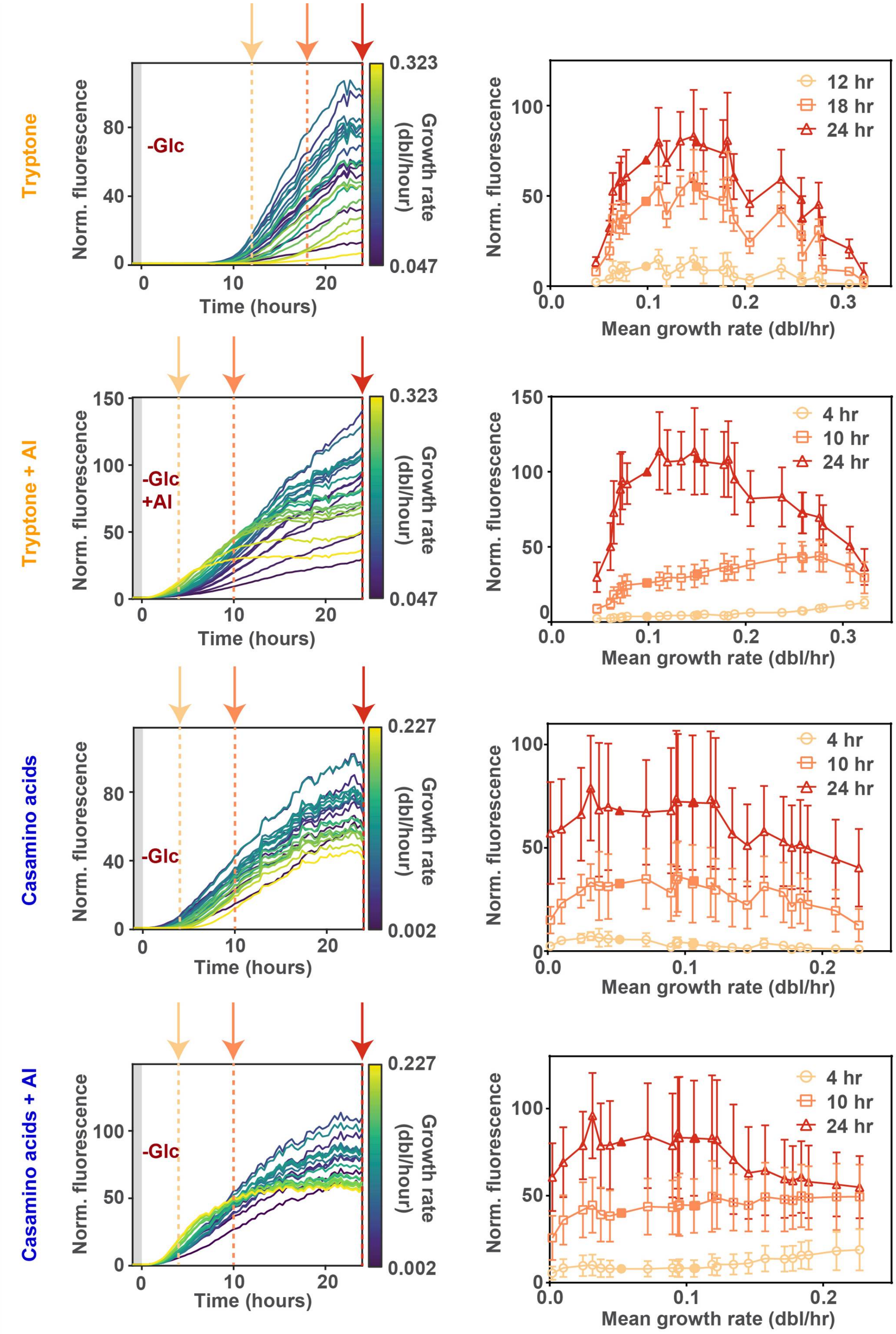


**Supplementary Figure 11 Time evolution of QS response distribution in various conditions.** Distributions of QS response at various time points in tryptone, tryptone with 1 μM exogenous

AI, casamino acids, and casamino acids with 1 μM exogenous AI medium at 25°C (n = 6, from 3 independent experiments) (mean ± SD). Filled shapes indicate the 2 chamber types excluded from this analysis due to partial formation of cell bilayer.

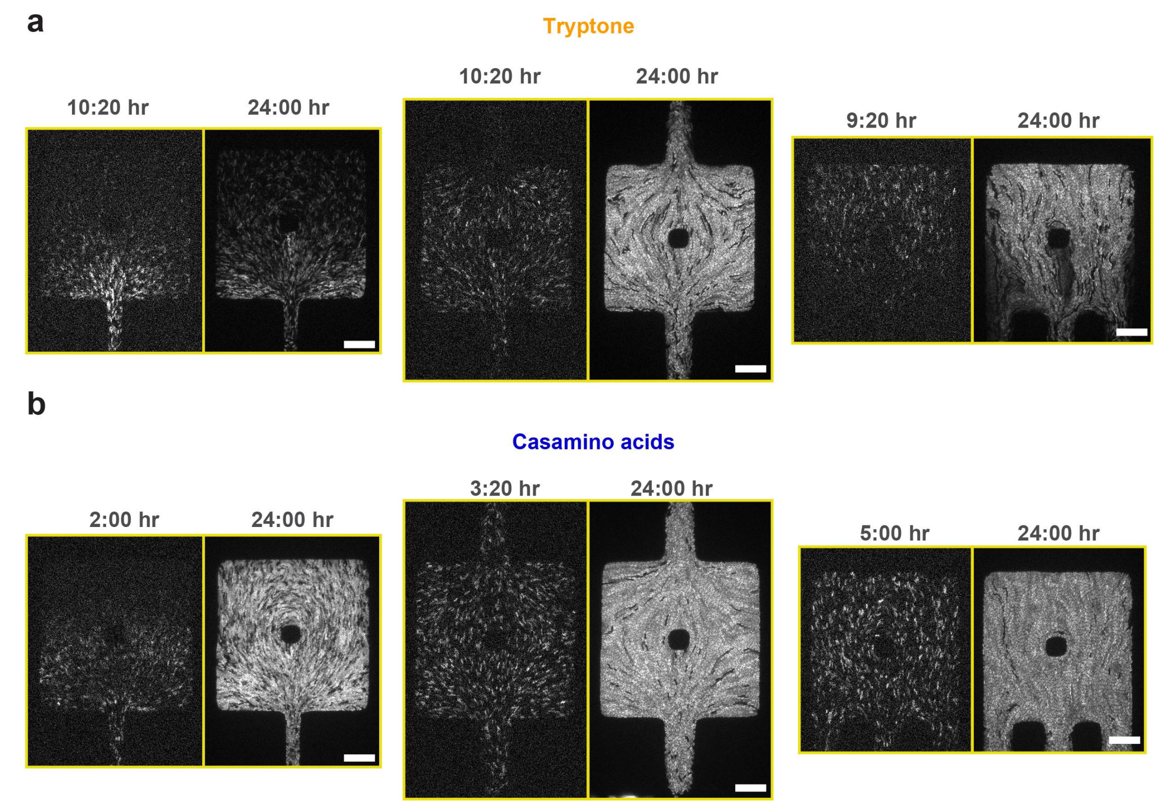


**Supplementary Figure 12 QS spatial response at onset and 24$^{th}$ hour in tryptone and casamino acids media conditions. a,b,** Fluorescence micrographs of various chamber configurations in **(a)** tryptone and **(b)** casamino acids media conditions at 25°C at onset (left) and 24$^{th}$ hour (right) time points**.** Scale bar, 20 µm.

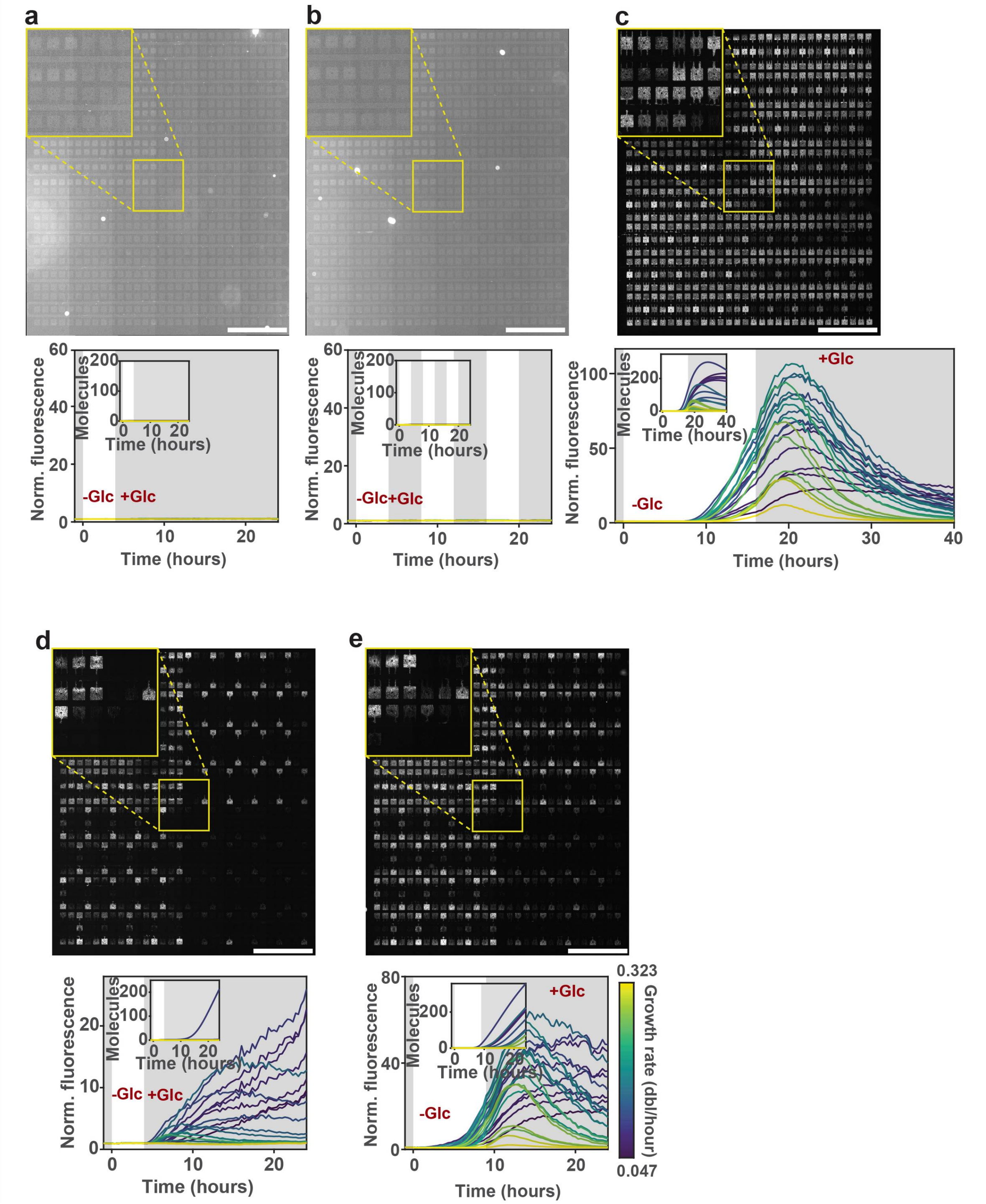


**Supplementary Figure 13 QS response within various transient environments. a-e,** Fluorescence micrographs and response dynamics of **(a)** 4 hour transient glucose (20 mM) removal, 4 hour, **(b)** 50% duty cycle pulses of glucose (20 mM) removal, **(c)** 16 hour transient glucose (20 mM) removal, **(d)** 4 hour transient glucose (10 mM) removal, and **(e)** 9 hour transient glucose (10 mM) removal (mean, n = 6, from 3 independent experiments). All conditions were in tryptone medium at 25°C. Inset in plot contains simulated dynamics. Inset in fluorescence micrograph contains magnified view of the region indicated, which contains all 24 chamber configurations. Scale bars, 1 mm.

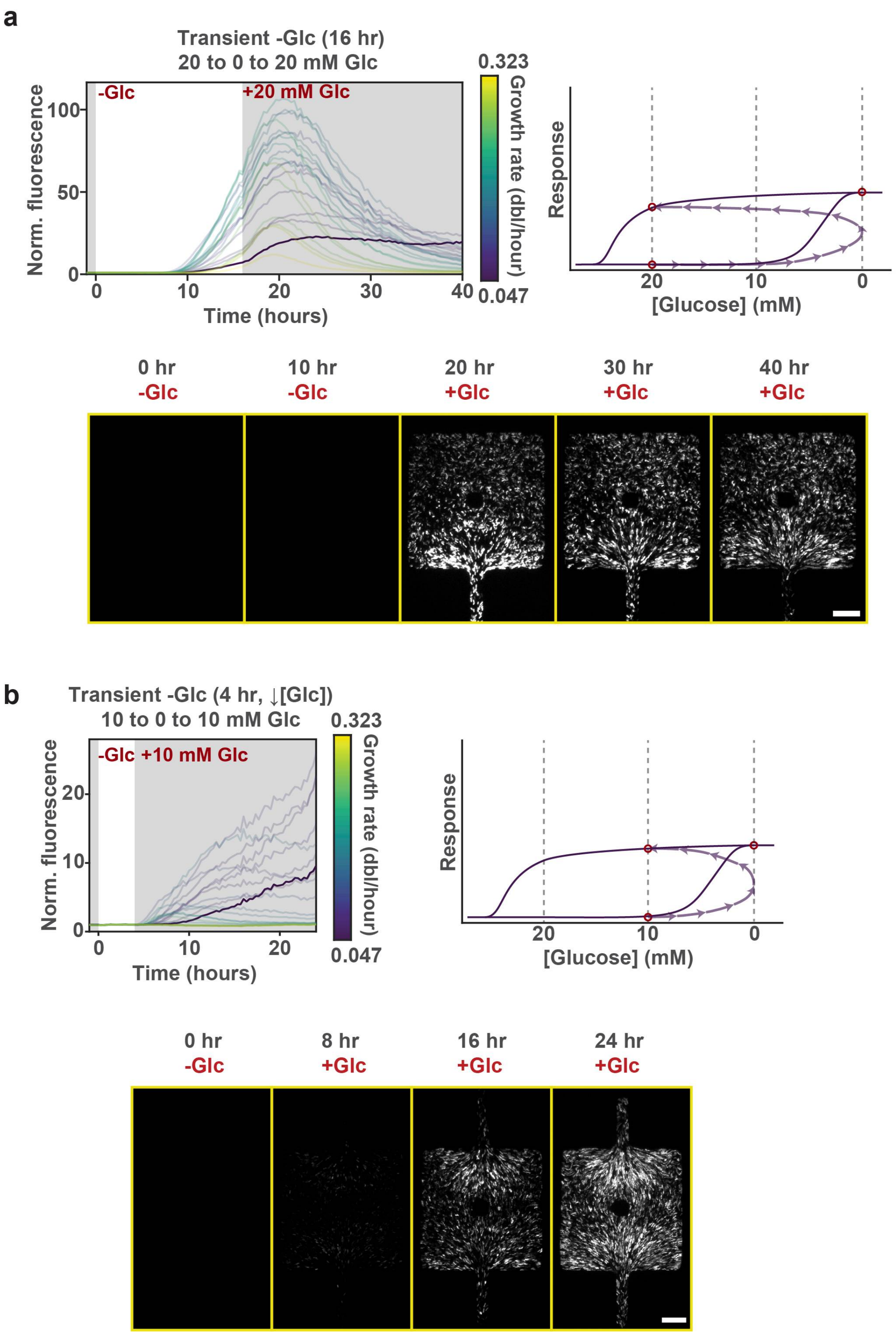


**Supplementary Figure 14 Bistability and hysteresis in transient environments. a,b**, Plot of response dynamics with accompanying diagram illustrating the hysteretic response for **(a)** 16

hour transient glucose (20 mM) removal and **(b)** 4 hour transient glucose (10 mM) removal. All conditions were in tryptone medium at 25°C (mean, n = 6, from 3 independent experiments). Filmstrip of representative chamber that maintained a stable high state of expression for each condition is shown. Scale bar, 20 μm.

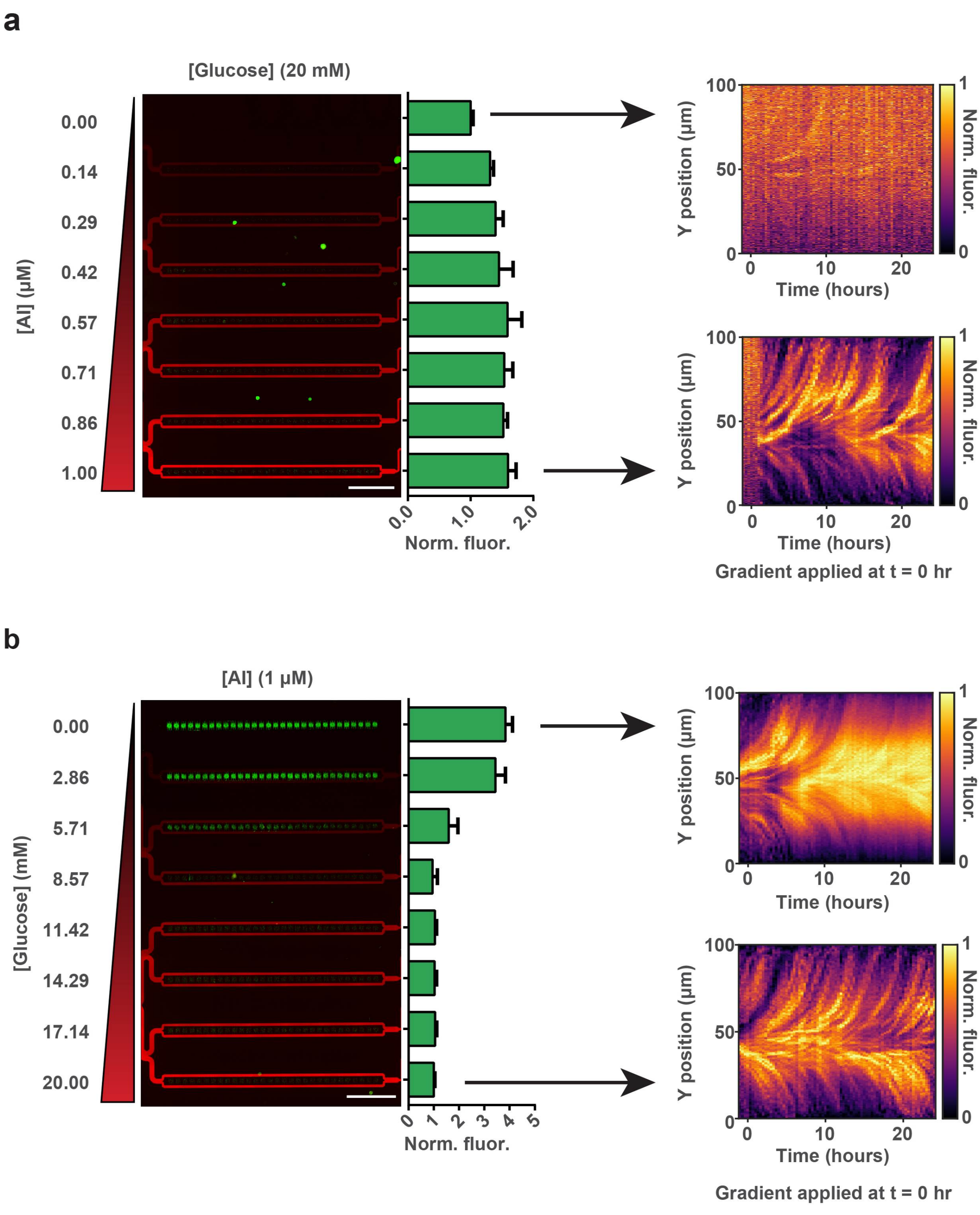


**Supplementary Figure 15 Simultaneous induction and repression elicits stably induced subpopulation. a**, Fluorescence micrograph and distribution of QS response in tryptone medium with 20 mM glucose and a 0-1 μM AI gradient at 25°C. Scale bar, 1 mm. Kymographs of the two extreme conditions are shown, with the gradient applied at time t = 0 hr. **b,** Fluorescence micrograph and distribution of QS response in tryptone medium with 1 μM exogenous AI and a 0-20 mM glucose gradient at 25°C. Scale bar, 1 mm. Kymographs of the two extreme conditions are shown, with the gradient applied at time t = 0 hr. Scale bar, 1 mm. (n=30) (mean ± SD)

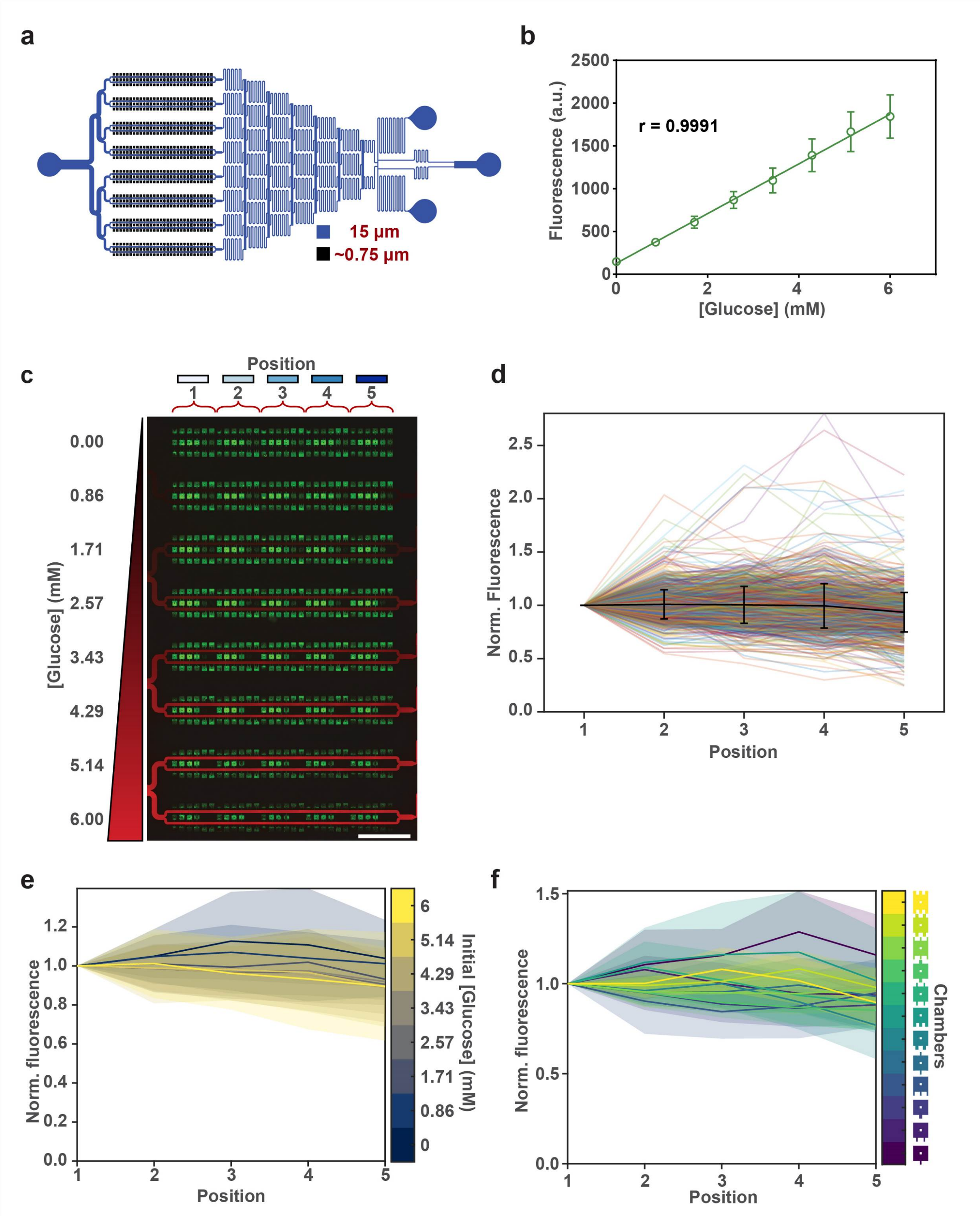


**Supplementary Figure 16 Characterization of gradient-generating microfluidic device for eliciting induced subpopulation. a,** Diagram of gradient-generating microfluidic device with 12 chamber configurations (short coupling channels). **b**, Dye fluorescence intensities within the 8 flow-through channels demonstrating linearity of the generated gradient (n=16, from 8 independent experiments). **c**, Fluorescence micrograph of gradient generating device, with positions of chamber clusters indicated and color-coded. **d**, Comparison of response intensity between the different positions, for all chamber configurations and dose concentrations. **e**, Comparison of response intensity between the different positions, for all chamber configurations across different dose concentrations. **f**, Comparison of response intensity between the different positions, for all dose concentrations across different chamber configuration. Scale bar, 1 mm. (mean ± SD)

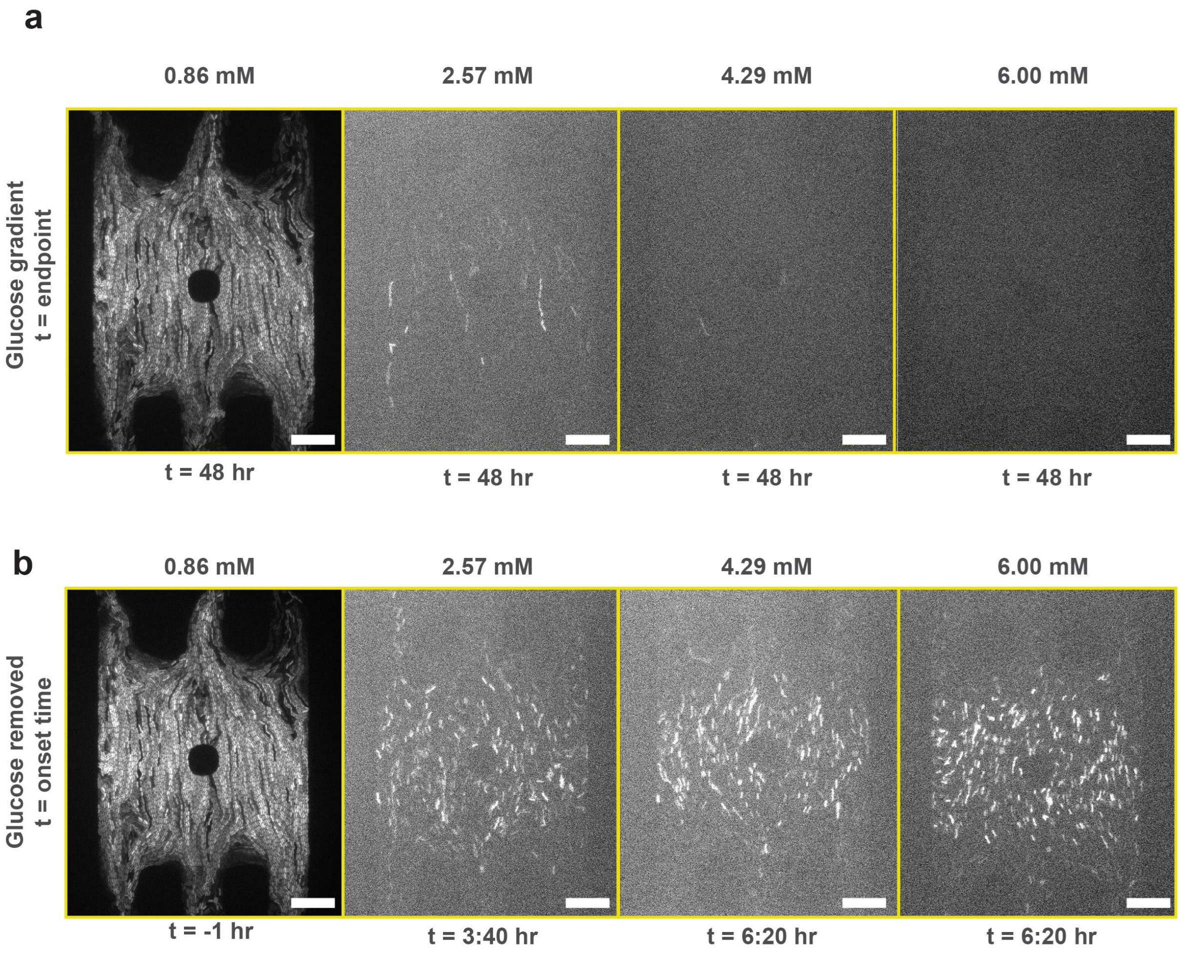


**Supplementary Figure 17 Comparison of onset time from different initial induction states.** **a,b,** Representative fluorescence micrographs of 4 dose conditions at **(a)** 48 hours after introduction of glucose gradient, and **(b)** time of onset after glucose gradient was removed. Scale bar, 20 µm.

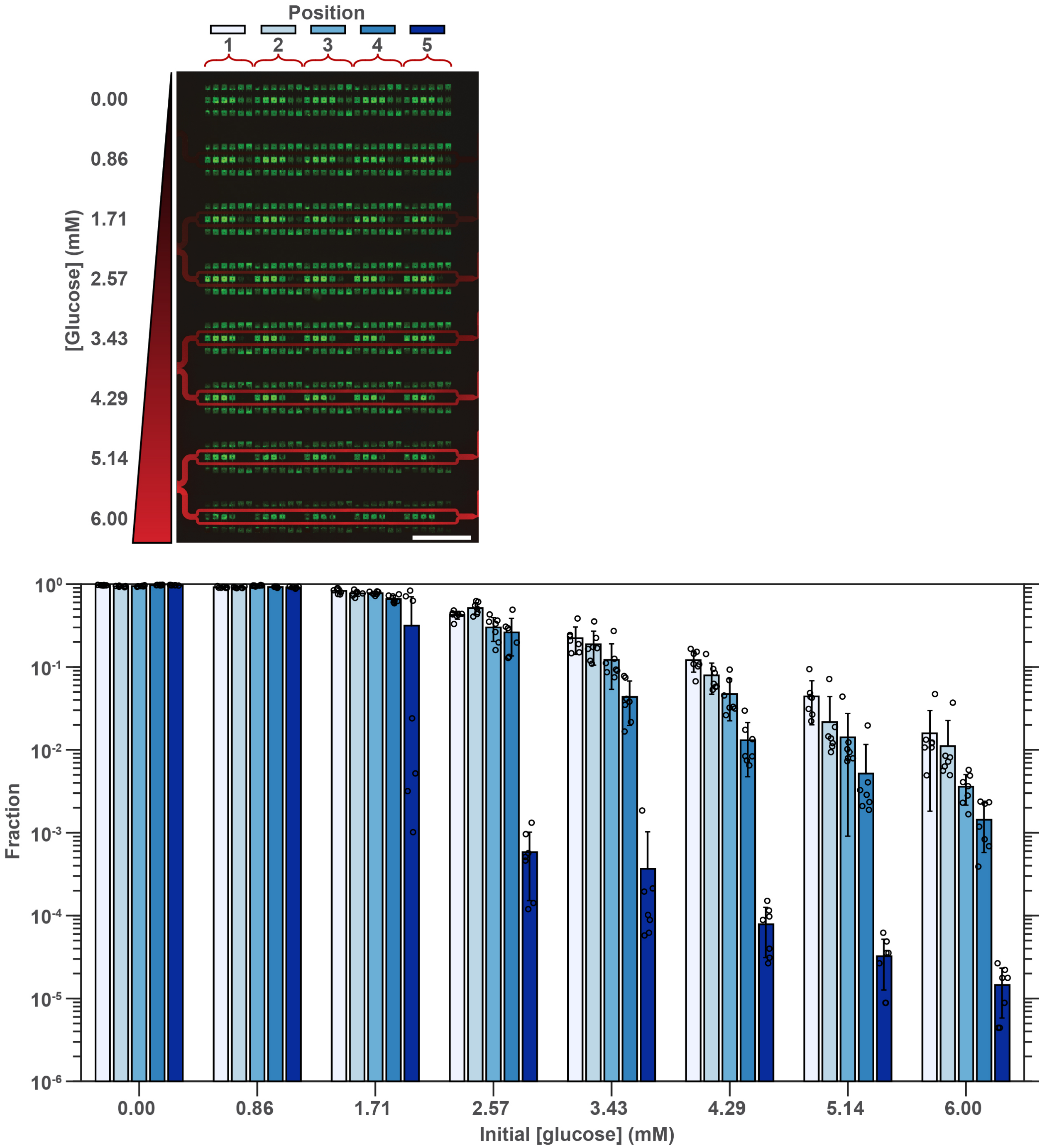


**Supplementary Figure 18 Fractions of induced cells.** The fraction of induced cells within various initial glucose concentrations and positions (n = 6, from 3 independent experiments) (mean ± SD). Scale bar, 1 mm.

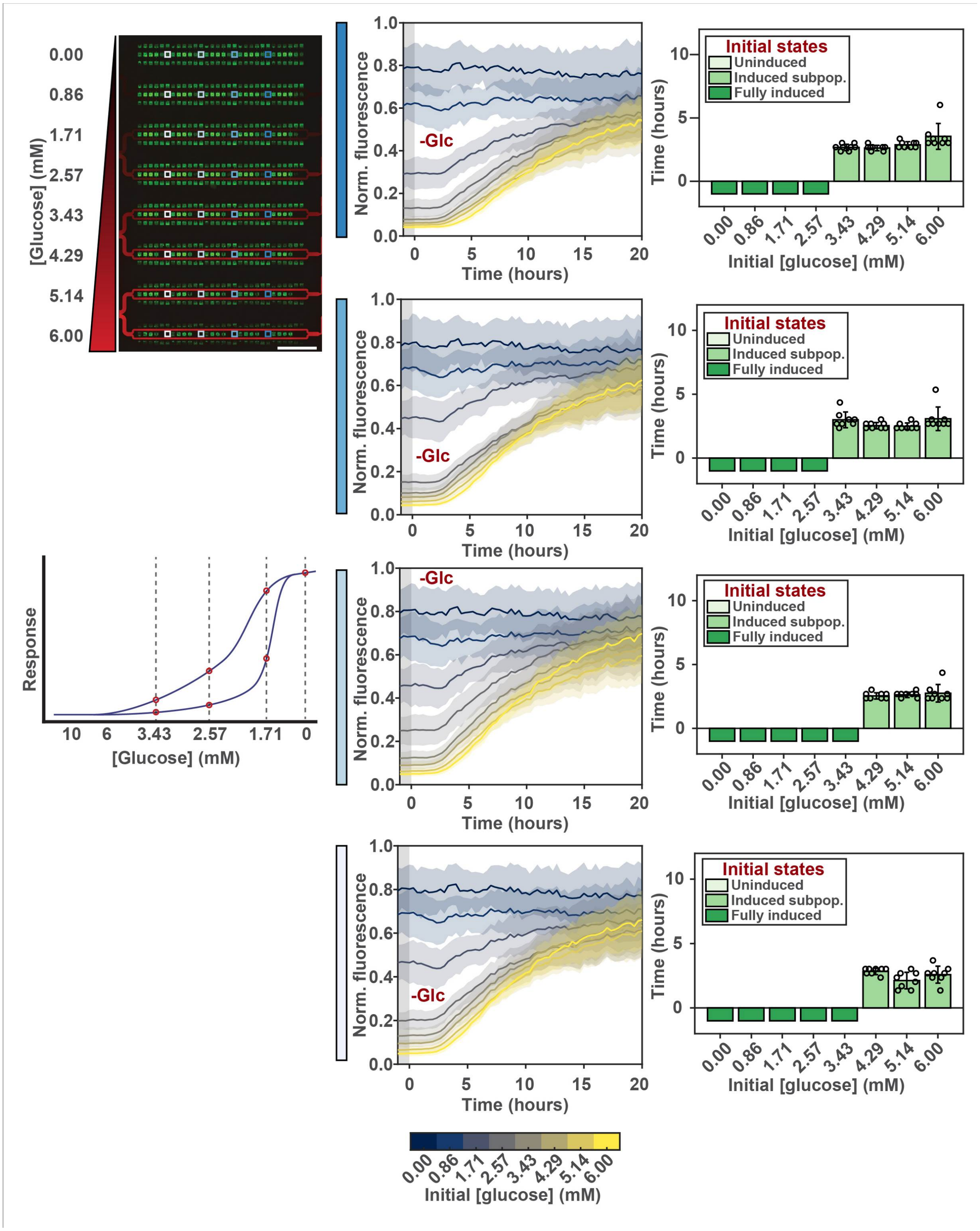


**Supplementary Figure 19 Response dynamics and onset time within various initial glucose concentrations and positions.** Plots of response dynamics with corresponding onset times at different positions and initial glucose concentrations (n = 8) (mean ± SD). Scale bar, 1 mm

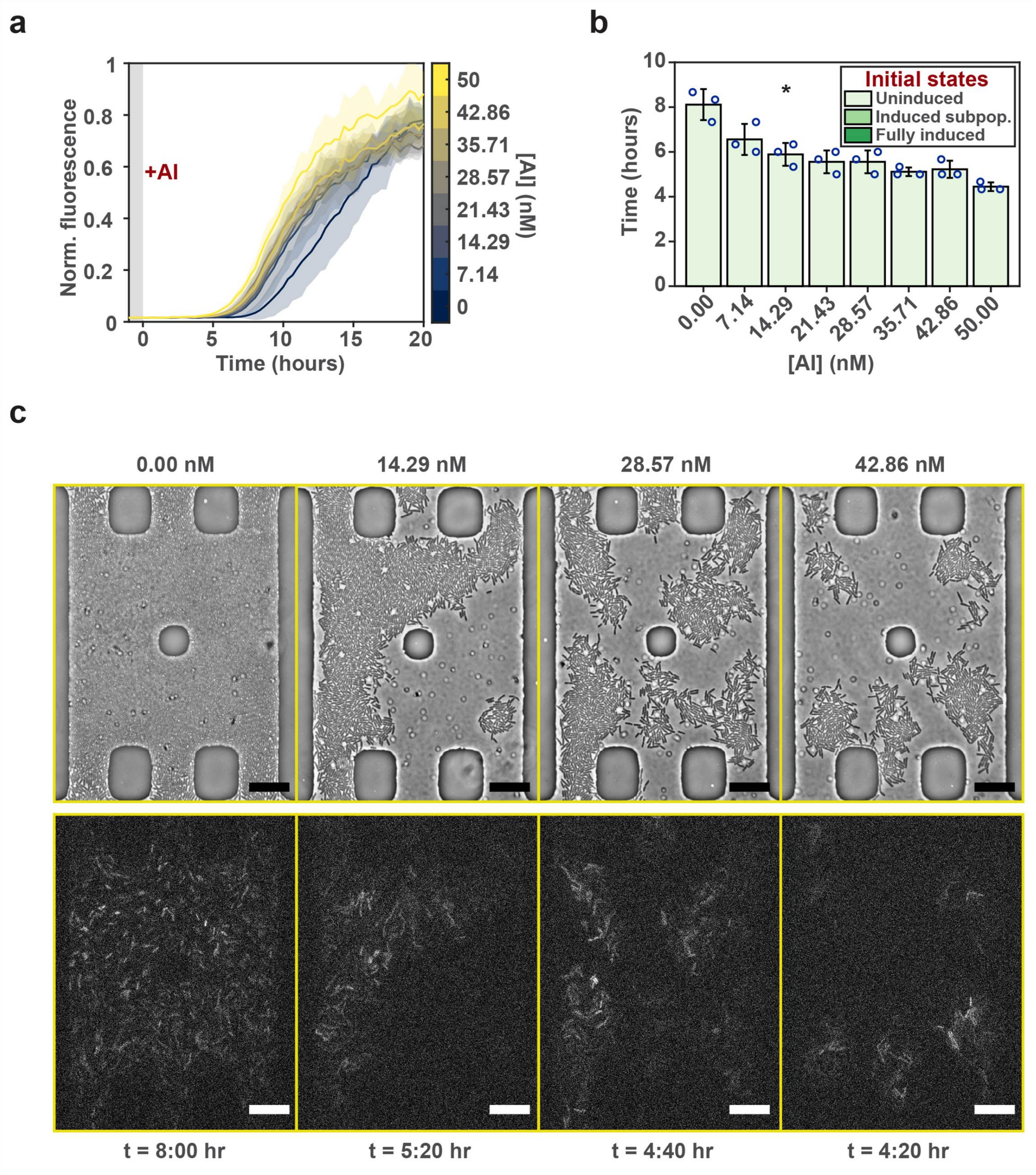


**Supplementary Figure 20 Low doses of AI is sufficient to reduce onset time. a,b,** Plots of **(a)** response dynamics and **(b)** onset time of QS response in tryptone medium with a 0-50 nM AI gradient at 25°C. (n = 3, *P<0.05, two tailed Student's t-test) (mean ± SD) **c,** Representative phase contrast and fluorescence micrographs of 4 dose conditions at time of onset after exogenous AI gradient was applied. Scale bar, 20 μm.

**Movie 1 Absence of QS response in tryptone medium with 20 mM glucose at 25°C for 24 hours.**

**Movie 2 Simulated dye diffusion.**

**Movie 3 Experimental dye diffusion.**

**Movie 4 Estimation of growth rate via GFP dilution.**

**Movie 5 QS response in tryptone medium at 25°C for 24 hours.**

**Movie 6 QS response in tryptone medium with 1 µM exogenous AI at 25°C for 24 hours.**

**Movie 7 QS response in alternating tryptone and casamino acids media with 1 µM exogenous AI at 25°C for 36 hours.**

**Movie 8 QS response in casamino acids medium at 25°C for 24 hours.**

**Movie 9 QS response in casamino acids medium with 1 µM exogenous AI at 25°C for 24 hours.**

**Movie 10 Absence of QS response in tryptone medium at 30°C for 48 hours.**

**Movie 11 QS response in casamino acids medium at 30°C for 24 hours.**

**Movie 12 QS response in tryptone medium with 1 mM IPTG (LuxR overexpression) at 30°C for 24 hours.**

**Movie 13 QS response in tryptone medium with 1 mM IPTG (LuxR overexpression) at 25°C for 24 hours.**

**Movie 14 QS response in tryptone medium with an initial 10 mM glucose at 25°C for 24 hours**

**Movie 15 Absence of QS response in tryptone medium at 25°C for 24 hours with 4 hour transient glucose (20 mM) removal.**

**Movie 16 Absence of QS response in tryptone medium at 25°C for 24 hours with 4 hour, 50% duty cycle pulses of glucose (20 mM) removal.**

**Movie 17 QS response in tryptone medium at 25°C for 40 hours with 16 hour transient glucose (20 mM) removal.**

**Movie 18 QS response in tryptone medium at 25°C for 24 hours with 4 hour transient glucose (10 mM) removal.**

**Movie 19 QS response in tryptone medium at 25°C for 24 hours with 9 hour transient glucose (10 mM) removal.**

**Movie 20 Simultaneous induction with 1 µM exogenous AI and repression with 20 mM glucose elicits stably induced subpopulation.**

**Movie 21 Fully induced, partially induced, and uninduced cell populations stably maintained in tryptone medium with 0.86, 2.57, 4.29, and 6 mM glucose at 25°C for 48 hours.**

**Movie 22 Onset of QS response in initially fully induced, partially induced, and fully uninduced cell populations in tryptone medium at 25°C for 20 hours.**

**Movie 23 Movie 22 with different dynamic range.**

**Movie 24 QS response in tryptone medium with 0, 14.29, 28.57, and 42.86 nM exogenous AI at 25°C for 24 hours.**

**Movie 25 Movie 24 with different dynamic range.**

**Movie 26 Phase contrast of movie 24.**
